# Cannabivarin and Tetrahydrocannabivarin Modulate Nociception via Vanilloid Channels and Cannabinoid-Like Receptors in *Caenorhabditis elegans*

**DOI:** 10.1101/2025.08.07.669105

**Authors:** Nasim Rahmani, Jesus D. Castaño, Francis Beaudry

**Author notes:** Corresponding author: Francis Beaudry, Ph.D., Professor of Analytical Pharmacology, Canada Research Chair in metrology of bioactive molecules and target discovery, Département de Biomédecine Vétérinaire, Faculté de Médecine Vétérinaire, Université de Montréal, 3200 Sicotte, Saint-Hyacinthe, QC, Canada J2S 2M2.

## Abstract

Cannabis has attracted growing interest for its therapeutic potential, especially in pain management. This study explores the antinociceptive effects of two promising non-psychoactive cannabinoids, cannabivarin (CBV) and tetrahydrocannabivarin (THCV), using *Caenorhabditis elegans* (*C. elegans*), a nematode model that expresses homologs of mammalian cannabinoid and vanilloid receptors. Thermotaxis assays were employed to quantify the antinociceptive effects of CBV and THCV in *C. elegans*. Wild-type animals were exposed to increasing concentrations of each compound to establish dose–response relationships. To investigate potential molecular targets, additional experiments were performed using mutant strains deficient in vanilloid receptor homologs (OCR-2 and OSM-9) and cannabinoid receptor homologs (NPR-19 and NPR-32). Mass spectrometry-based proteomics combined with network biology analyses were used to identify the biological pathways associated with drug response. Results confirmed that both compounds elicit dose-dependent antinociceptive effects. Mutant analyses support the involvement of vanilloid and cannabinoid signaling pathways in mediating these responses. These findings highlight the potential of CBV and THCV as non-psychoactive analgesic agents and support further research into their mechanisms of action and translational relevance for mammalian pain management.

## Introduction

Pain is a complex sensory and emotional experience involving nociceptive pathways that detect harmful stimuli and transmit signals to the central nervous system (Raja et al., 2020). Pain detection and transmission are mediated by nociceptors, which respond to harmful mechanical, thermal, or chemical stimuli. Nociceptive signals travel from the periphery to the central nervous system, where they may be interpreted as pain (St John Smith, 2018, Dubin and Patapoutian, 2010, Woolf and Ma, 2007, Smith and Lewin, 2009, Brodal, 2010, Hucho and Levine, 2007, Loeser and Melzack, 1999). While nociception is a neural process that can occur without conscious awareness, pain is a subjective and multidimensional experience involving sensory, emotional, and cognitive components (Baliki and Apkarian, 2015, Garland, 2012). Understanding these pathways is essential for developing more effective analgesic strategies (Tracey and Mantyh, 2007).

Cannabinoids have shown promise as analgesic agents through interactions with cannabinoid (CB1 and CB2) and vanilloid receptors (Ross, 2003, Vučković et al., 2018). Δ9-tetrahydrocannabinol (THC) derived from *Cannabis sativa* is the most studied phytocannabinoid with therapeutic properties; however, its psychoactivity limits its usage (Aggarwal et al., 2009). Despite significant progress in elucidating the pharmacological properties of major cannabinoids such as Δ9-tetrahydrocannabinol (THC) and cannabidiol (CBD), our understanding of many other phytocannabinoids and their metabolites remains limited (Hanuš et al., 2016). Among these lesser-known compounds, cannabivarin (CBV) and tetrahydrocannabivarin (THCV) have recently attracted attention due to their structural similarity to THC and their potential therapeutic properties (Li et al., 2024b). Preliminary studies suggest that these cannabinoids may interact with the endocannabinoid and vanilloid systems in distinct ways, possibly exerting unique effects on nociception, inflammation, and neuroprotection (Sampson, 2021, Stone et al., 2020). However, the precise molecular mechanisms through which CBV and THCV act remain poorly defined. Critical questions persist regarding their binding affinities to cannabinoid receptors (CB1 and CB2), potential interactions with non-canonical targets such as TRP channels or GPR55, and their downstream signaling pathways. Moreover, the functional consequences of their receptor engagement, particularly *in vivo*, are still largely unexplored. Further investigation is essential to clarify their pharmacodynamics and therapeutic potential, and to understand how they differ from or complement more extensively studied cannabinoids.

This study investigates the antinociceptive effects of CBV and THCV using *Caenorhabditis elegans* (*C. elegans*), a genetically tractable model organism that expresses mammals’ homologs of cannabinoid and vanilloid receptors (Boujenoui et al., 2024, Abdollahi et al., 2024, Nkambeu et al., 2020). We hypothesized that CBV and THCV exert antinociceptive effects in *C. elegans* through interactions with homologous cannabinoid receptors (NPR-19 and NPR-32) and vanilloid receptors (OSM-9 and OCR-2). Our objectives were to (1) characterize their exposure– response relationships using heat avoidance assays, (2) identify the relevant receptor targets, and (3) elucidate the proteins and pathways involved in their antinociceptive activity through mass spectrometry-based proteomics. These findings may offer new insights into cannabinoid-based therapeutics and the molecular underpinnings of nociception.

## Materials and methods

### Chemicals and reagents

All chemicals and reagents used in this study were purchased from Fisher Scientific (Fair Lawn, NJ, USA) or Millipore Sigma (St. Louis, MO, USA). CBV and THCV were obtained from Toronto Research Chemicals (North York, ON, Canada).

### *C.elegans* strains

The *C.elegans* N2 (Bristol) strain was used as the reference strain. The mutant strains included npr-19 (ok2068), npr-32 (ok2541), ocr-2 (ak47), and osm-9 (ky10), which are associated with cannabinoid and vanilloid receptor homlogs. All strains were obtained from the Caenorhabditis Genetics Center (CGC) at the University of Minnesota (Minneapolis, MN, USA). Following standard protocols, nematodes were cultured and maintained on nematode growth medium (NGM) agar (Brenner, 1974, Margie et al., 2013). Routine maintenance was carried out at 22°C in a refrigerated incubator, and all experimental procedures were performed at room temperature (∼22°C) unless stated otherwise.

### *C. elegans* pharmacological manipulations

CBV and THCV stock solutions (1 mg/mL) were prepared and diluted in Type 1 ultrapure water to a final concentration of 25 μM. The solutions were thoroughly vortexed to ensure homogeneity. Serial dilutions were performed to obtain further concentrations of 10 μM, 5 μM, and 1 μM in Type 1 ultrapure water. Nematodes were collected and washed following the protocol described by Margie et al. (Margie et al., 2013). After 72 hours of feeding and development on NGM plates (92 × 16 mm Petri dishes), nematodes were removed from food and exposed to CBV or THCV solutions. A 7 mL aliquot of CBV or THCV solution was added to create a thin solution film (2–3 mm), partially absorbed by the NGM, allowing nematodes to remain suspended in the solution. The nematodes were exposed to CBV or THCV for 60 minutes, then isolated and washed three times with S Basal before behavioral assessments. To evaluate residual effects, nematodes were transferred to NGM plates free of CBV or THCV for a 6-hour latency period following exposure before behavioral testing.

### Thermal avoidance assays

The thermal avoidance assay used in this study was adapted from the previously established four-quadrant method (Margie et al., 2013) and has been extensively utilized in prior research (Boujenoui et al., 2024, Lahaise et al., 2024, Abdollahi et al., 2024, Nkambeu et al., 2021, Nkambeu et al., 2020). Experiments were conducted in 92 × 16 mm Petri dishes divided into four quadrants. A central circular region (1 cm in diameter) was designated for nematode placement but was excluded from analysis. Two quadrants (A and D) served as heat stimulus zones, while the remaining quadrants (B and C) functioned as controls. To immobilize nematodes, all quadrants were treated with 0.5 M sodium azide. A noxious heat stimulus was applied using an electronically heated metal tip (0.8 mm in diameter), positioned 2 mm above the nematode growth medium (NGM) agar. This setup generated a radial temperature gradient, maintaining a temperature range of 32–35°C, as confirmed using an infrared thermometer. The stimulus temperature was selected based on prior studies (Wittenburg and Baumeister, 1999). Before testing, nematodes were collected and washed according to Margie et al. (Margie et al., 2013). Groups of 100–300 nematodes were placed in the center of the marked Petri dish. After a 30-minute stimulation period, plates were stored at 4°C for at least one hour. A digital stereomicroscope was used to count the number of nematodes in each quadrant, excluding those remaining within the inner circle. The thermal avoidance index (TI) and the percentage of animal avoidance behavior were calculated to assess nocifensive responses to noxious heat.

### Sample preparation for proteomics

Nematodes, either untreated or exposed to CBV or THCV, were collected in liquid medium, centrifuged at 1,000 × g for 10 minutes, and thoroughly washed with S Basal. The washed nematodes were then resuspended in phosphate-buffered saline (PBS) containing 137 mM NaCl, 2.7 mM KCl, 10 mM Na HPO, and 1.8 mM KH PO, supplemented with 1% (v/v) Triton X-100 and a cOmplete™ protease inhibitor cocktail (Roche Diagnostic Canada, Laval, QC, Canada). The suspensions were transferred into reinforced 1.5 mL homogenizer tubes containing 25 mg of 500 μm glass beads. Homogenization was carried out using a Bead Mill Homogenizer (Fisher Scientific) with five bursts of 60 seconds at a speed of 5 m/s. The resulting homogenates were centrifuged at 12,000 × g for 10 minutes, and protein concentrations were determined using the Bradford assay.

For protein extraction, approximately 100 µg of protein was precipitated using ice-cold acetone (1:5, v/v). The protein pellets were dissolved in 100 µL of 50 mM TRIS-HCl buffer (pH 8.0) and subjected to vigorous vortexing (2,800 rpm) followed by sonication to enhance solubility. To denature proteins, the solution was heated at 120°C for 10 minutes in a reaction block and subsequently cooled for 15 minutes. Proteins were then reduced with 20 mM dithiothreitol (DTT) at 90°C for 15 minutes, followed by alkylation with 40 mM iodoacetamide (IAA) in the dark at room temperature for 30 minutes. Proteolytic digestion was performed by adding 5 µg of proteomics-grade trypsin to each sample, with incubation at 37°C for 24 hours. The reaction was terminated by adding 10 µL (∼10% v/v) of a 1% trifluoroacetic acid (TFA) solution. The final samples were centrifuged at 12,000 × g for 10 minutes, and 100 µL of the supernatant was transferred into HPLC vials for mass spectrometry analysis.

### Proteomics analysis

Online chromatographic separation was performed using a Thermo Scientific Vanquish Neo UHPLC system (San Jose, CA, USA) configured in trap-and-elute mode. The setup included a Thermo Scientific PepMap Neo 5 μm C18 300 μm × 5 mm trap cartridge and a Thermo Scientific PepMap Neo C18 2 μm × 75 μm × 500 mm nano column. A 2 μL sample (∼2 μg of digested proteins) was loaded onto the trap column and desalted with solvent A [0.1% formic acid in water] at a 20 μL/min flow rate for 0.5 minutes. Peptide elution was carried out using a linear gradient of 5% to 35% solvent B [80% acetonitrile, 20% water, 0.1% formic acid] over 135 minutes, followed by a 2-step column wash at 50% solvent B for 10 minutes and 80% for 10 minutes. The flow rate was maintained at 200 nL/min. Afterwards, the column was equilibrated to the initial solvent composition (5% solvent B) by running 10 column volumes. Data acquisition was conducted on a Thermo Scientific Q Exactive Plus Orbitrap Mass Spectrometer (San Jose, CA, USA) equipped with a Nanospray Flex ion source. Data collection was performed in positive ion mode with a nanospray voltage of 2.2 kV, and the ion transfer tube was maintained at 220°C. The mass spectrometer was operated in data-dependent acquisition (DDA) mode, utilizing a TOP-12 acquisition strategy. A high-resolution MS^1^ survey scan (m/z 375– 1200) was acquired at a resolution of 70,000 (FWHM), with an automatic gain control (AGC) target value of 1 × 10 and a maximum injection time of 100 ms. The 12 most intense precursor ions exceeding an intensity threshold of 1 × 10 were selected for fragmentation via higher-energy collisional dissociation (HCD) at a normalized collision energy (NCE) of 27. MS^2^ spectra were acquired at a resolution of 17,500 (FWHM) using an AGC target value of 2 × 10 and a maximum injection time of 100 ms.

### Protein quantification

Protein identification and quantification were performed using FragPipe (version 22.0) with the MSFragger search engine. Data processing was conducted using default settings unless specified otherwise. The raw MS^1^ and MS^2^ spectra were analyzed against the *C. elegans* reference proteome, obtained from UniProt (taxon identifier 6239) in FASTA format. Database search parameters included a precursor mass and fragment mass tolerance of 20 ppm. Trypsin was set as the digestion enzyme, allowing up to two missed cleavages. Fixed modifications included carbamidomethylation of cysteine, while variable modifications accounted for methionine oxidation, N-terminal acetylation, and phosphorylation. A minimum peptide length of seven amino acids was required for inclusion in the analysis. False discovery rate (FDR) filtering was performed using Percolator, with an FDR threshold of 1%. Relative quantification was conducted using a label-free approach, integrating peak intensities with ionQuant default options within FragPipe, and analyzing MaxLFQ intensity with FragPipe Analyst.

### Bioinformatics

Differential protein expression analysis was conducted using datasets generated by FragPipe (version 22.0) and its online tool FragPipe-Analyst Extracted parameters included the log abundance ratio (experimental group vs. N2 control), corresponding p-values, p-adjusted values, and protein accession identifiers. Functional enrichment analysis was performed using ClusterProfiler, incorporating Gene Ontology (GO) biological process (BP) annotation, and Reactome Pathways (Wu et al., 2021). UMAP plots were produced using the uwot package (Melville, 2025) in the software R (version 4.4.1).

### Statistical analysis

Behavioral data were analyzed using the nonparametric Kruskal–Wallis test, followed by Dunn’s post-hoc test for multiple comparisons. Statistical significance was set at p ≤ 0.05. All statistical analyses and figure generation were conducted using GraphPad Prism (version 10.5.0, GraphPad Software, Inc., La Jolla, CA, USA).

## Results and Discussion

### Thermal avoidance assessment

We first assessed whether CBV or THCV exposure affected baseline locomotion or quadrant preference in *C. elegans* under non-stimulus conditions. As shown in Figure 1, both wild-type (N2) and mutant *C. elegans* strains exhibited no significant quadrant bias at 22°C, and drug exposure did not alter their spatial distribution 30 minutes post-placement compared to the untreated group. These results confirmed there is no experimental bias without heat stimulus. Figure 2 illustrates the dose-dependent effects of cannabivarin (CBV) on thermal avoidance behavior in *C. elegans*. A 60-minute exposure to CBV led to a significant hindrance of nocifensive responses to noxious heat, consistent with an antinociceptive effect. Notably, this reduction in heat avoidance behavior was more pronounced at higher CBV concentrations (10 µM and 25 µM), indicating a concentration-dependent modulation of nociceptive signaling. Furthermore, the antinociceptive effect of CBV was not transient; behavioral assays conducted up to six hours after exposure, revealed a sustained decrease of nocifensive responses to noxious heat. These findings suggest that CBV elicits a prolonged effect on nociceptive circuits, potentially through lasting receptor engagement or downstream signaling modulation. The persistence of this effect warrants further investigation into the pharmacological properties of CBV and its potential to induce long-term alterations in sensory processing.

**Figure 1.**
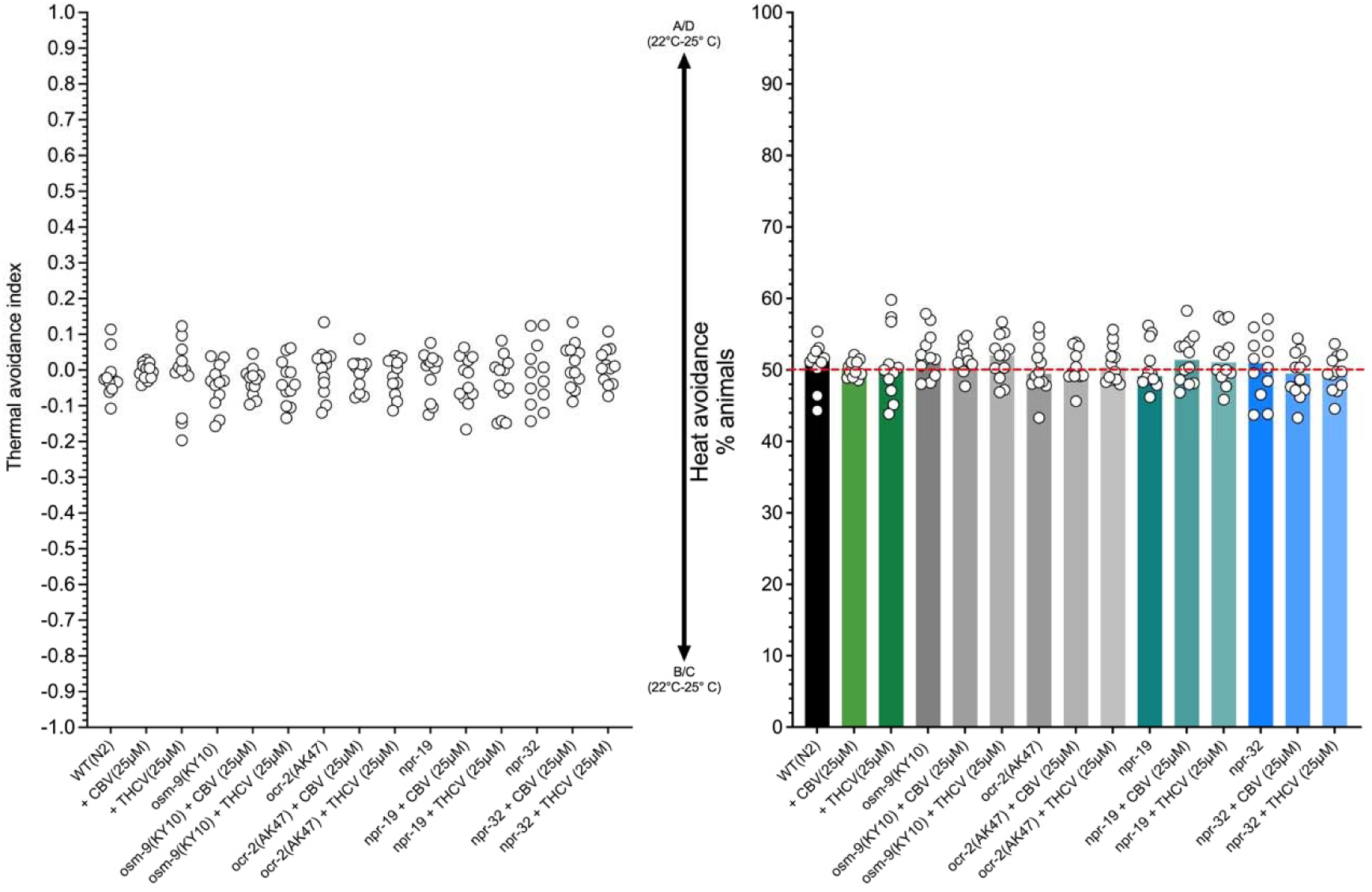
Comparison of mobility and quadrant bias between wild-type (N2) and selected mutant *C. elegans* strains on quadrant plates maintained at a constant temperature of 22°C under no-stimulus conditions (negative control). The tested genotypes exhibit no significant quadrant preference, irrespective of the presence or absence of CBV or THCV. ****p < 0.0001, *** p < 0.001, ** p < 0,01, * p < 0.05 (non-parametric Kruskal-Wallis analysis followed by Dunn’s multiple comparisons test).

**Figure 2.**
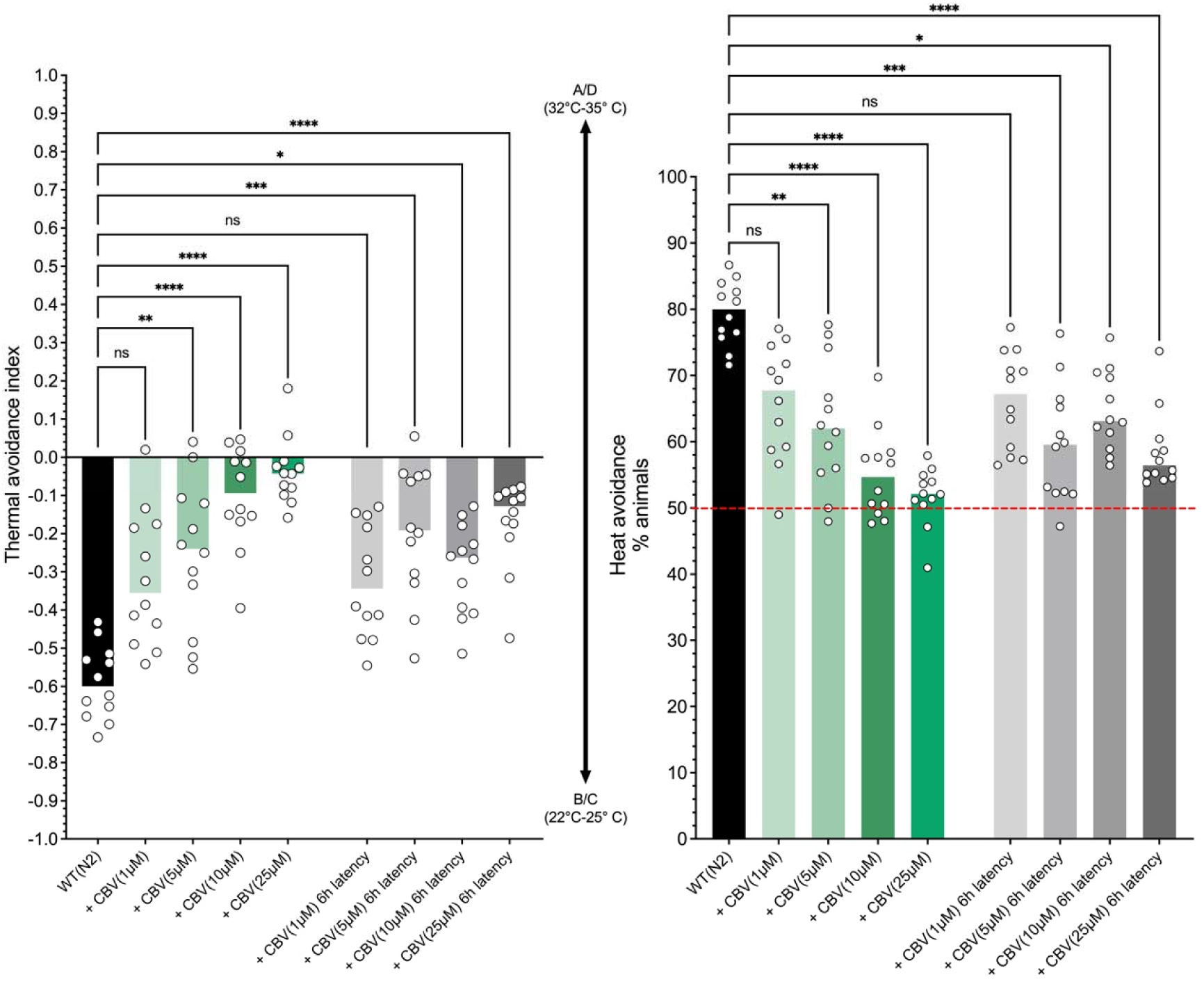
Evaluation of the pharmacological effects of CBV on thermal avoidance behavior in *C. elegans*. Nematodes were exposed to CBV for 60 minutes before undergoing behavioral assays. Data points represent individual values, with medians shown, derived from at least 12 independent experiments per experimental group. CBV demonstrates dose-dependent effects, significantly impairing thermal avoidance responses in *C. elegans*. Residual antinociceptive effects are marked 6 hours post-exposure. ****p < 0.0001, *** p < 0.001, ** p < 0,01, * p < 0.05 (non-parametric Kruskal-Wallis analysis followed by Dunn’s multiple comparisons test).

THCV also caused dose-dependent inhibition of nocifensive behaviour in *C. elegans*, as shown in Figure 3. Similar to CBV, higher concentrations of THCV (10 µM and 25 µM) produced a more significant reduction in thermal avoidance, supporting its antinociceptive potential. However, unlike CBV, the effects of THCV did not last after a 6-hour washout period, with nocifensive responses returning to near-baseline levels. This transient effect suggests that THCV may undergo rapid metabolic degradation or clearance in *C. elegans*, or that its receptor binding is brief and characterised by low affinity or short-lived interactions. The lack of residual antinociceptive activity contrasts with the sustained effects seen with CBV and indicates possible differences in receptor affinity, pharmacokinetics, or downstream signalling stability between the two compounds. The prolonged effects of CBV may reflect extended receptor engagement or slower metabolic turnover, highlighting important pharmacodynamic differences that merit further molecular and pharmacological investigation.

**Figure 3.**
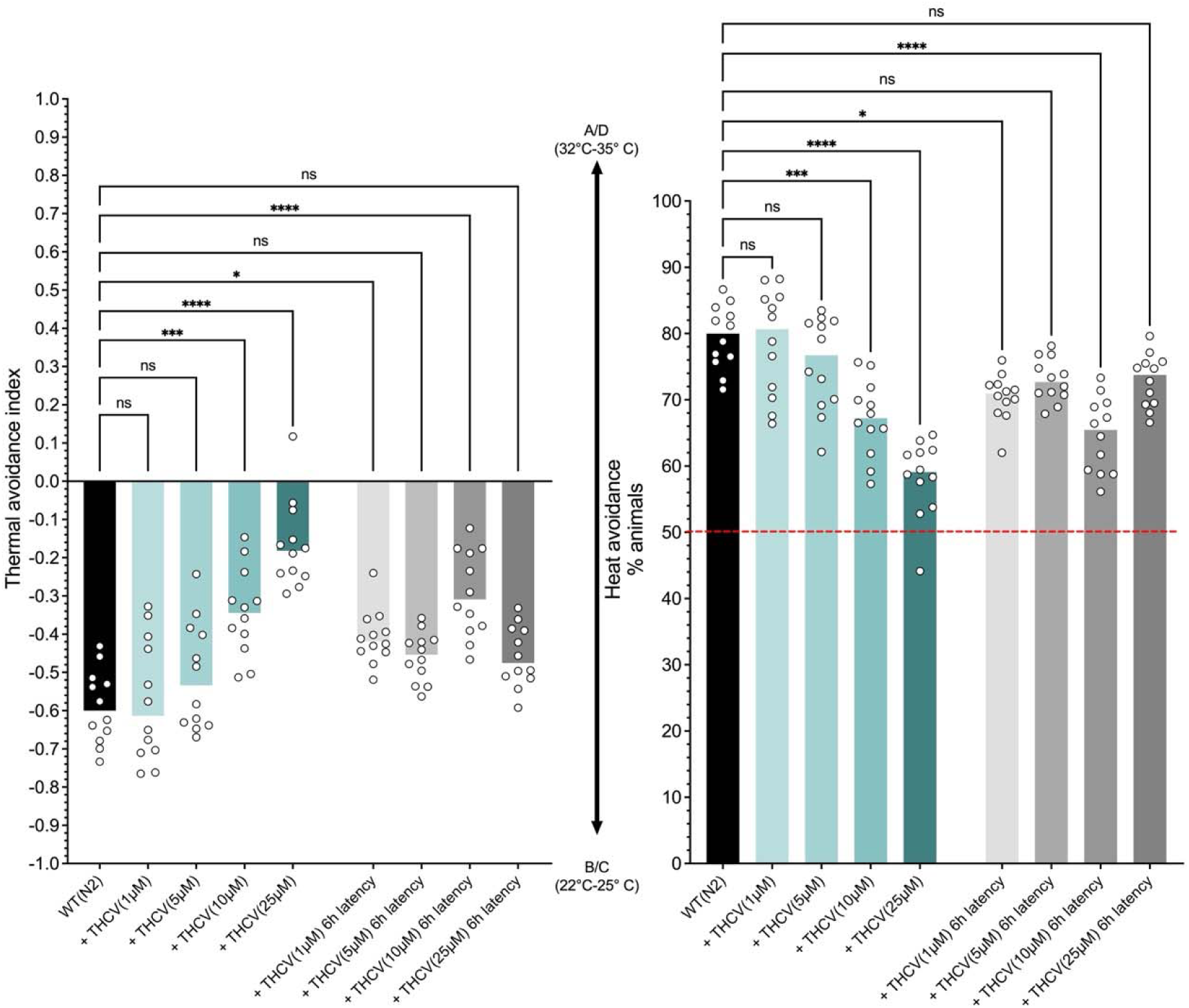
Assessment of THCV’s pharmacological effect on thermal avoidance behavior in *C. elegans*. Nematodes were treated with THCV for 60 minutes prior to behavioral testing. Data represent individual values alongside medians, based on at least 12 independent experiments per group. THCV exhibits dose-dependent activity, notably reducing thermal avoidance responses in *C. elegans*. Significant residual antinociceptive effects persist up to 6 hours after exposure. ****p < 0.0001, *** p < 0.001, ** p < 0,01, * p < 0.05 (non-parametric Kruskal-Wallis analysis followed by Dunn’s multiple comparisons test).

### Identification of CBV and THCV Targets

To identify the receptor targets responsible for CBV and THCV’s effects, we assessed thermal avoidance behavior in receptor-deficient *C. elegans* strains. We used strains lacking functional vanilloid receptors (*ocr-2*, *osm-9*) and cannabinoid receptors (*npr-19*, *npr-32*). Mutant nematodes were exposed to 25 μM CBV or THCV for 60 minutes before behavioral assessments. As shown in Figure 4, the antinociceptive effects of CBV were not significantly altered in *osm-9* and *npr-32* mutants, suggesting that its action is primarily mediated through the vanilloid receptor OSM-9 and the cannabinoid-like receptor NPR-32. This pattern is consistent with previous reports indicating that CBV has low affinity for CB1 and CB2 receptors but exhibit moderate activity at TRP ion channels, including TRPV1 (Sampson, 2021). In contrast, THCV’s effects were impaired in all four tested mutants (*ocr-2*, *osm-9*, *npr-19*, and *npr-32*), suggesting broader receptor involvement or functional redundancy among nociceptive pathways. However, the limited sensitivity of the behavioral assay may have hindered the detection of subtle or modest effects on nociceptive behavior. Previous studies have also shown that THCV, a structural analogue of THC, exhibits reduced binding affinity for CB1 and CB2 receptors due to its shorter aliphatic side chain and is known to modulate TRP channels, particularly TRPV1 (Pertwee, 2008, Hill et al., 2012, De Petrocellis et al., 2011). The absence of THCV-induced antinociceptive effects in *ocr-2*, *osm-9*, *npr-19*, and *npr-32* mutants, despite its clear activity in WT nematodes, suggests that these receptors are mediators of THCV’s action. This supports the conclusion that THCV requires the functional presence of both TRP channel orthologs (OSM-9 and OCR-2) and cannabinoid-like receptors (NPR-19 and NPR-32) to exert its antinociceptive effects (Pinho-Ribeiro et al., 2017, Nkambeu et al., 2021, Nkambeu et al., 2020, Salem et al., 2022). These findings highlight the complexity of nociceptive signaling in *C. elegans*, and suggest that THCV interacts with multiple, non-redundant molecular targets to modulate thermal avoidance behaviour (Abdollahi et al., 2024, Oakes et al., 2019).

**Figure 4.**
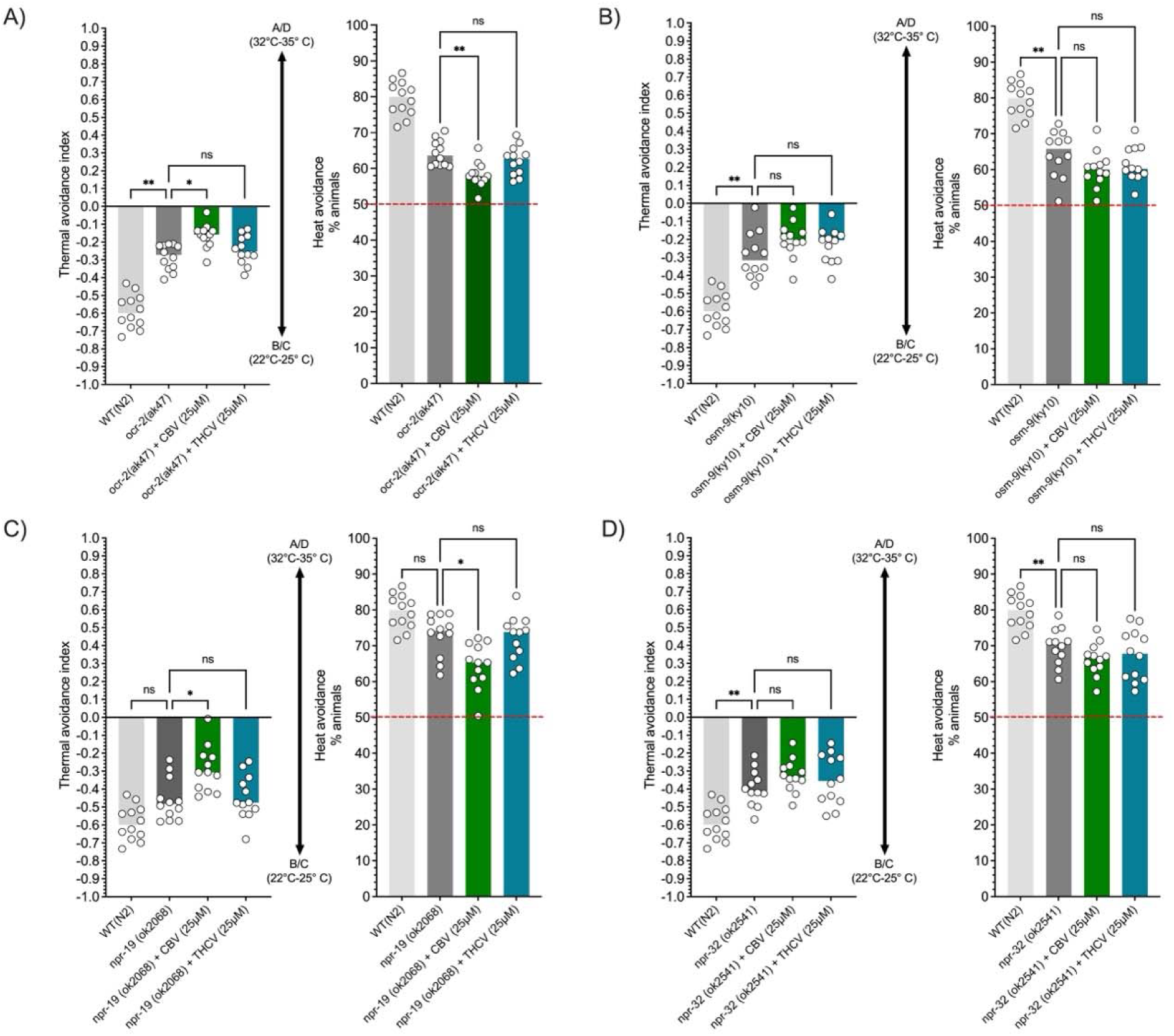
Identification of vanilloid receptor orthologs involved in CBV- and THCV-induced antinociceptive effects. Data points represent individual values with medians indicated, derived from at least 12 independent experiments per group. Tested mutants included ocr-2 (A), osm-9 (B), npr-19 (C), and npr-32 (D). CBV’s antinociceptive effects appear to be mediated by the vanilloid receptor OSM-9 and the cannabinoid receptor NPR-32. THCV’s antinociceptive effects may involve OCR-2, OSM-9, NPR-19, and NPR-32, implicating both vanilloid and cannabinoid receptor pathways. ****p < 0.0001, *** p < 0.001, ** p < 0,01, * p < 0.05 (non-parametric Kruskal-Wallis analysis followed by Dunn’s multiple comparisons test).

### Proteomic and Bioinformatic Investigations

Label-free quantitative proteomics was performed using mass spectrometry to assess global protein expression changes following a 1-hour exposure to CBV (25 µM) and THCV (25 µM). Comparative analysis identified differentially expressed proteins (DEPs) relative to untreated controls. Functional enrichment analysis and network biology approaches were applied to investigate biological processes and pathways associated with these DEPs. The UMAP visualization of protein abundance, shown in Figure 5, reveals protein landscapes between the control, CBV-, and THCV-treated groups, indicating substantial proteomic shifts following exposure to phytocannabinoids. In the control group, several protein clusters (circled in red) exhibit unique abundance signatures, including high- and low-abundance regions that are markedly altered or diminished in the CBV and THCV conditions. This suggests that both compounds broadly remodel the proteomic landscape. CBV treatment appears to disrupt these distinct clusters more uniformly, leading to a more homogenized distribution of protein expression, which may reflect its sustained and widespread molecular effects observed in prior analyses. In contrast, THCV treatment maintains some regional structure while still redistributing specific proteins, consistent with its more transient and targeted mode of action. The disappearance or reorganization of the circled clusters in the treated groups highlights specific protein subsets that may be directly involved in the antinociceptive response or downstream regulatory pathways, warranting further targeted validation.

**Figure 5.**
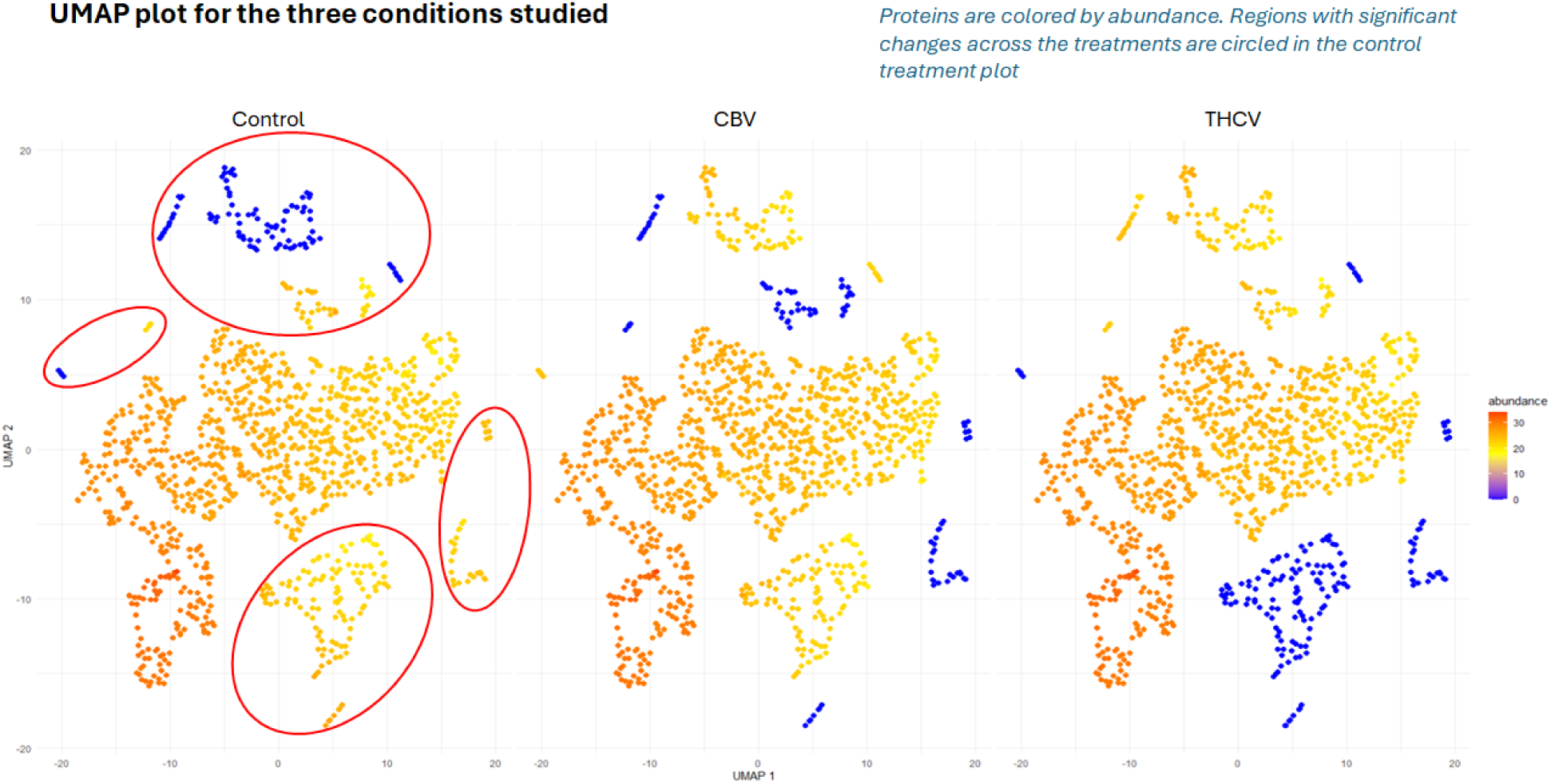
UMAP visualization of protein abundance across treatment conditions. Each point represents a protein, colored by abundance (purple = low, yellow = high). UMAP projections reveal distinct clustering patterns in control, CBV-, and THCV-treated samples. Red circles highlight regions in the control condition where protein abundance patterns are significantly altered in response to CBV or THCV exposure. The loss or redistribution of these clusters in treated groups suggests compound-specific proteomic remodeling, consistent with the sustained effects of CBV and the transient, targeted activity of THCV.

The gene set enrichment analysis (GSEA) comparing the biological processes affected by CBV and THCV reveals distinct patterns of pathway activation and suppression. As shown in Figure 6-A, both CBV and THCV primarily activated biological processes rather than suppressing them. CBV treatment led to the upregulation of pathways associated with negative regulation of signaling, particularly those involved in the response to stimuli, signal transduction, and cell communication. These findings suggest a dampening of excitatory signaling pathways, which may underlie CBV’s sustained antinociceptive effects observed behaviorally. CBV also uniquely activated processes such as cell migration, tissue development, and chromatin remodeling, suggesting broader impacts on cellular function that may indirectly influence nociceptive signaling by modulating neuronal plasticity, gene expression, or neural circuit development. Importantly, while THCV shared some overlapping activation with CBV, including regulation of developmental growth and metabolic processes, it demonstrated a narrower range of affected pathways. This divergence is further visualized in the hierarchical clustering shown in Figure 6-B, where CBV-associated processes clustered more distinctly and extensively than those of THCV, indicating broader molecular engagement. Notably, THCV induced fewer significant changes in signaling regulation and showed limited impact on pathways such as chromatin remodeling. On the suppression axis, THCV modestly downregulated small molecule biosynthesis and meiotic processes, indicating transient and possibly compensatory cellular responses. These differences support the behavioral data, where THCV produced shorter-lived antinociceptive effects. The broader and more sustained pathway engagement by CBV may reflect its prolonged interaction with receptor targets or differential intracellular signaling dynamics. These results are coherent with known pain modulation mechanisms in mammals, where the negative regulation of signal transduction and cell communication is associated with reduced neuronal excitability and diminished pain perception (Ossipov et al., 2010). The activation of pathways related to tissue development and chromatin remodeling by CBV may reflect longer-term neuroplastic changes, similar to those observed with sustained analgesic effects in mammalian models (Song et al., 2024, Denk and McMahon, 2012). In contrast, THCV’s more limited and transient impact on these pathways aligns with its reported short-acting effects and weaker modulation of central pain circuits.

**Figure 6.**
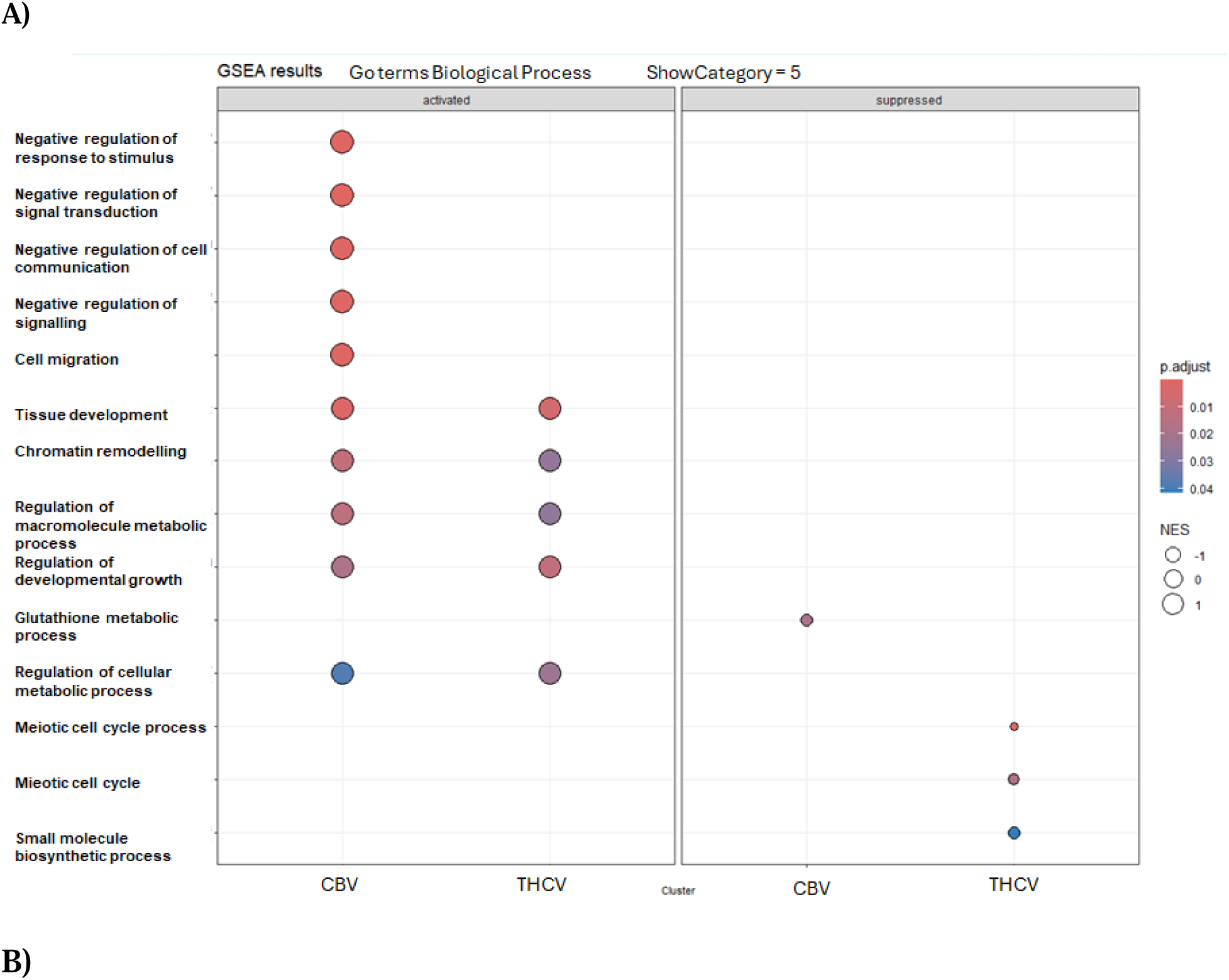
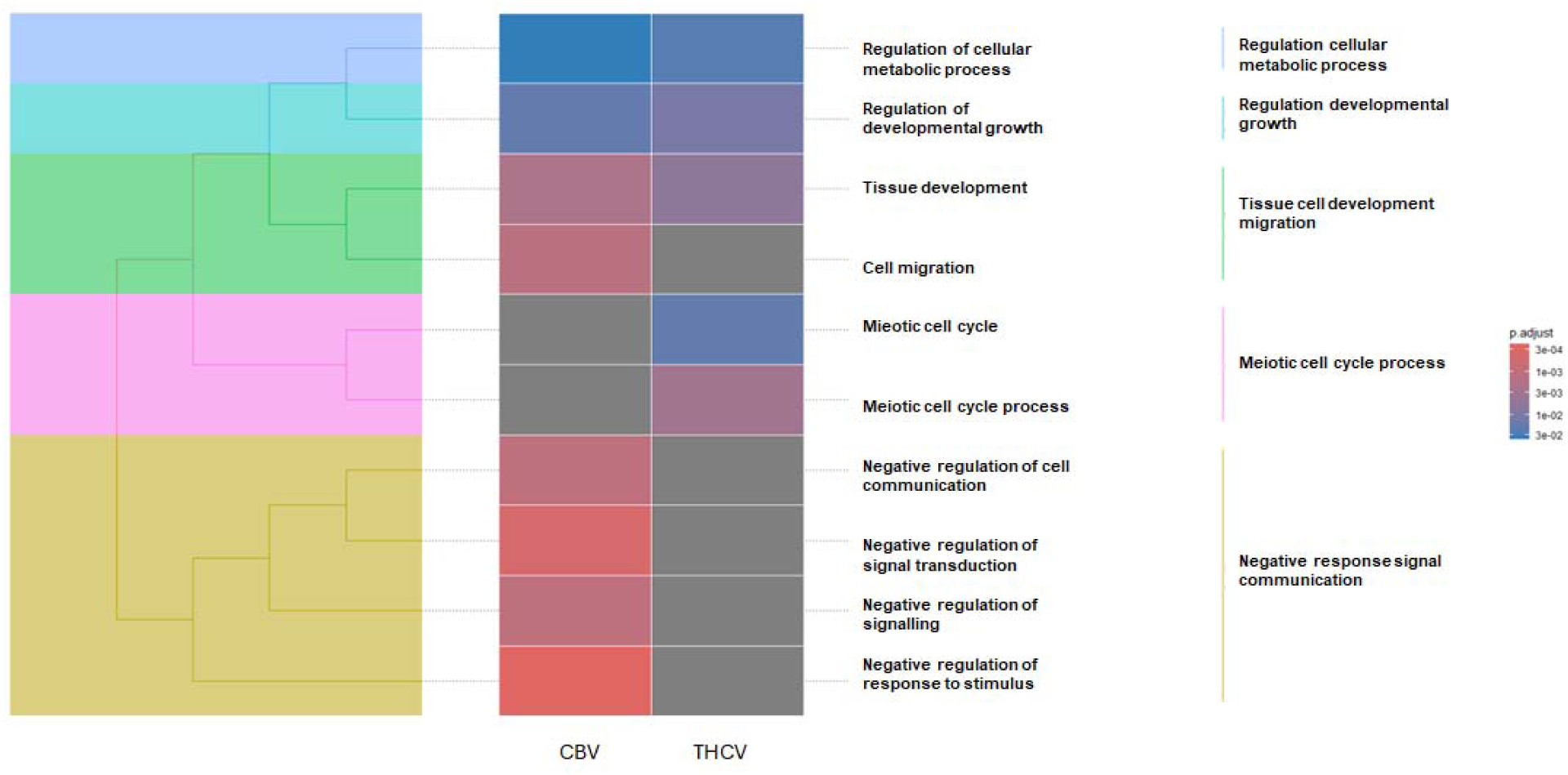
**(A).** GSEA dot plot of enriched GO Biological Processes in CBV- and THCV-treated animals. Biological processes significantly enriched following CBV and THCV exposure are shown, separated into activated (left panel) and suppressed (right panel) categories. Dot color indicates adjusted p-value, while size represents normalized enrichment score (NES). CBV treatment broadly activated pathways associated with negative regulation of stimulus response, signal transduction, cell communication, tissue development, and chromatin remodeling, consistent with sustained proteomic and behavioral effects. THCV induced a narrower range of activated processes, with modest enrichment in metabolic and developmental pathways. Suppression was more prominent in THCV-treated samples, particularly affecting meiotic and biosynthetic processes. These distinct pathway profiles suggest compound-specific mechanisms of antinociceptive action. **(B).** Tree plot of enriched Gene Ontology biological processes (GO-BP) in CBV- and THCV-treated *C. elegans*. The plot displays semantically grouped GO terms based on their similarity, revealing distinct biological process enrichments for CBV and THCV. CBV triggered broader activation of categories related to signaling regulation, cell communication, and tissue development, while THCV showed more limited and selective enrichment. Color scale represents adjusted p-values

The Reactome pathway analysis highlights distinct signaling responses induced by CBV and THCV, underscoring differences in their mechanisms of action. As shown in Figure 7-A, CBV strongly activated pathways related to gene expression, including RNA polymerase II transcription and global transcriptional regulation. This suggests that CBV may exert broad genomic effects, potentially contributing to its sustained antinociceptive activity observed in behavioral assays. In addition, CBV noticeably induced signaling pathways such as WNT and general signal transduction processes, both of which are associated with neuroplasticity and cell communication in mammalian systems (Rosso and Inestrosa, 2013, Narvaes and Furini, 2022, Tang et al., 2022, Li et al., 2024a). These pathway-specific differences are also reflected in the hierarchical clustering shown in Figure 7-B, where CBV-targeted processes form distinct branches, indicating stronger and more coordinated molecular engagement than THCV. WNT signaling plays an important role in the regulation of nociception and pain through its influence on neuronal development, plasticity, and inflammation (Zhou et al., 2022). This highly conserved pathway, originally known for its role in embryogenesis and cell fate determination, has emerged as a key modulator of pain pathways in both the central and peripheral nervous systems (Zhou et al., 2022, Zhao and Yang, 2018, Itokazu et al., 2014, Cao et al., 2024). Moreover, WNT pathway components have been shown to interact with TRP channels and opioid signaling, further linking WNT activity to nociceptive modulation (Rapetti-Mauss et al., 2020). As such, targeting WNT signaling represents a promising strategy for novel analgesic development, particularly in conditions involving neuropathic or inflammatory pain. CBV also upregulated pathways involved in mitotic cell cycle progression, M phase regulation, and drug metabolism (ADME), indicating a more extensive cellular and metabolic engagement. These findings may suggest broader modulation of cellular homeostasis and stress responses, which have been implicated in glial proliferation, synaptic remodeling, and neuroimmune crosstalk, processes that contribute to central sensitization and pain chronification (Grace et al., 2014, Ji et al., 2016, Inoue and Tsuda, 2009).

**Figure 7.**
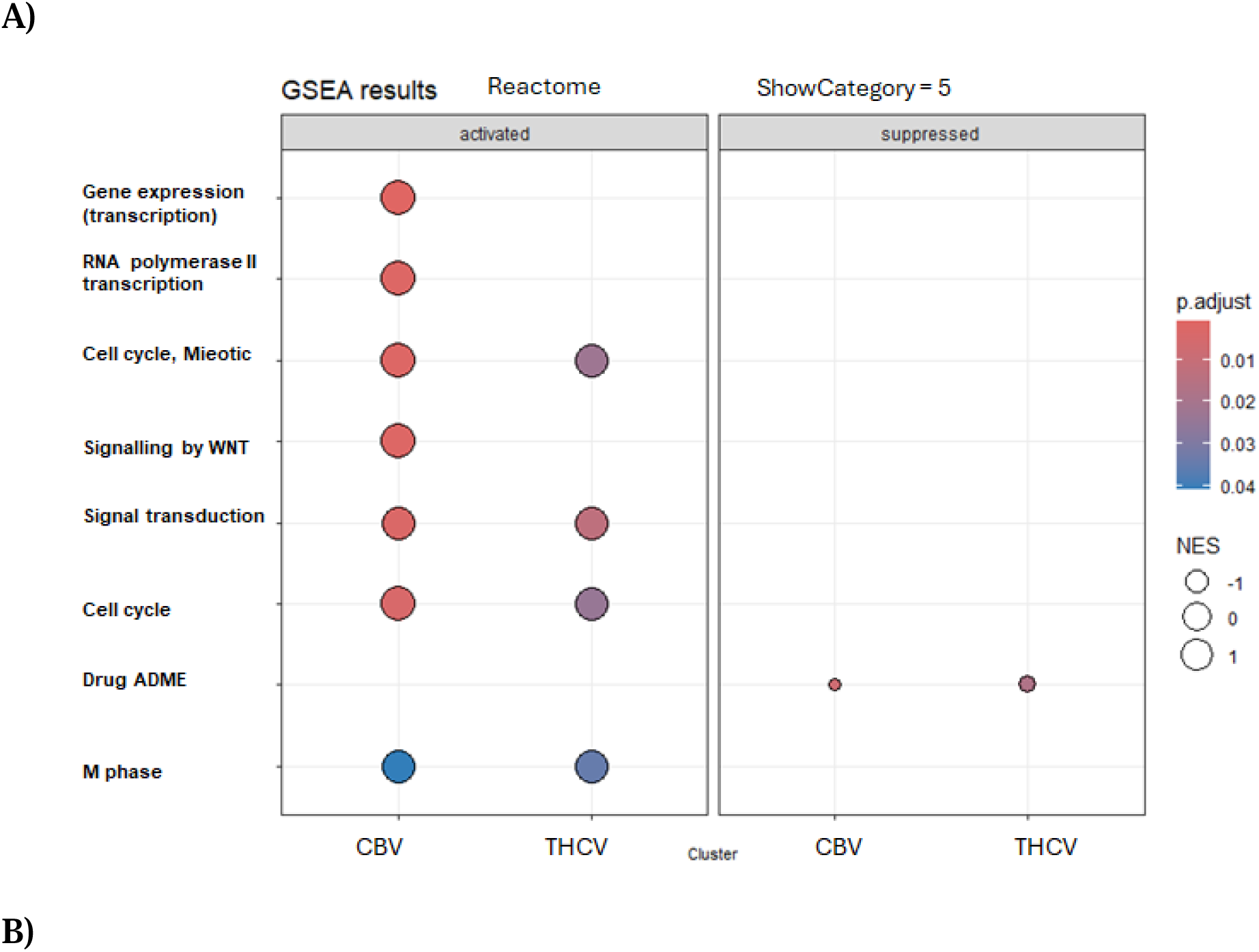
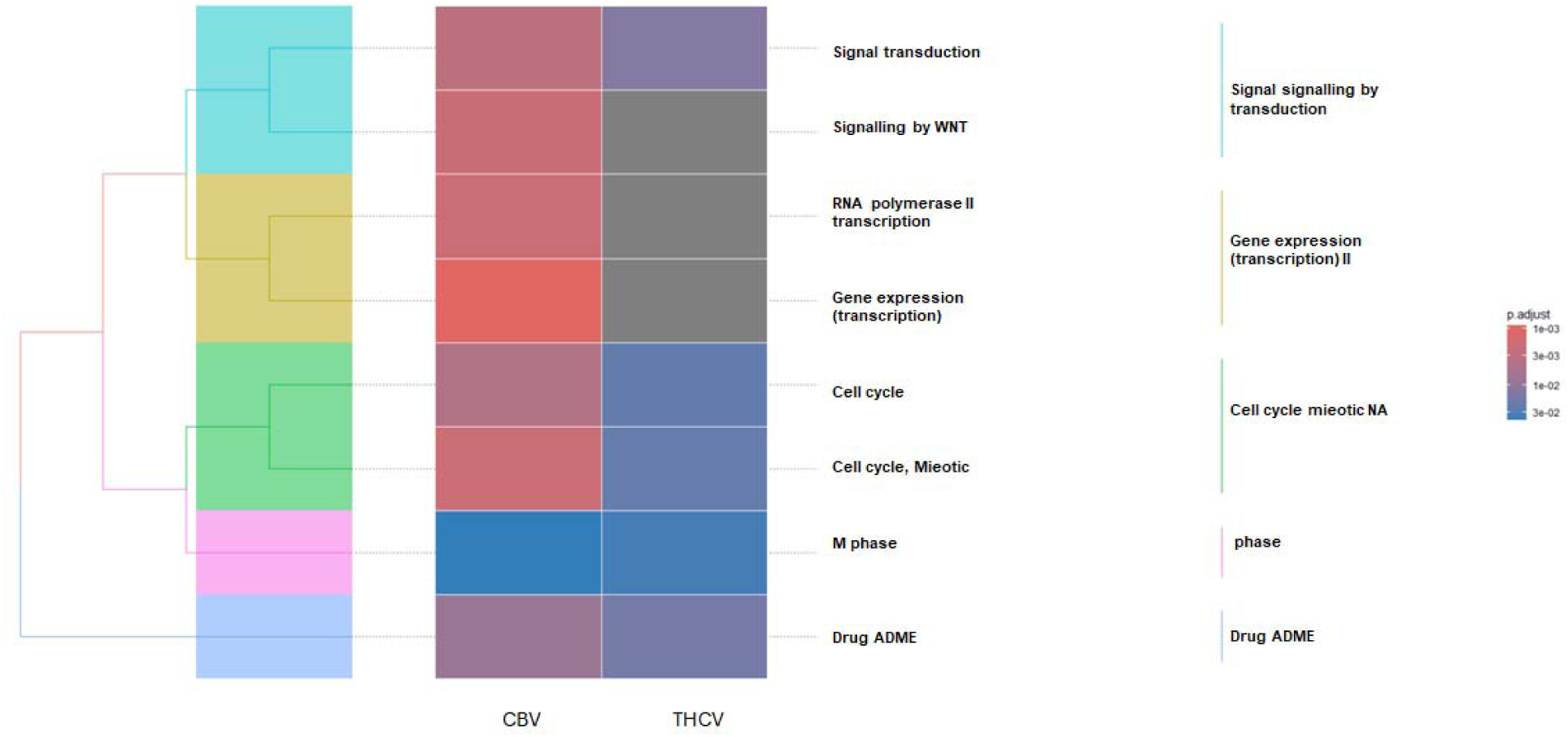
**(A).** Reactome pathway enrichment analysis of CBV- and THCV-treated *C. elegans*. Dot plot showing significantly enriched pathways in CBV and THCV conditions, separated into activated (left panel) and suppressed (right panel) categories. Dot size reflects normalized enrichment score (NES), and color indicates adjusted p-value. CBV robustly activated pathways involved in transcription (RNA polymerase II and global gene expression), WNT signaling, general signal transduction, mitotic cell cycle, and drug metabolism (ADME), suggesting widespread cellular engagement. THCV showed moderate enrichment in a subset of these pathways, with reduced breadth and intensity. Minimal suppression was observed for either compound. These results highlight distinct mechanistic profiles, with CBV engaging broader genomic and signaling programs linked to sustained antinociceptive effects. (**B).** Tree plot of enriched Reactome pathways in CBV- and THCV-treated *C. elegans*. Reactome pathway enrichment results are grouped by semantic similarity. CBV induced coordinated activation of transcriptional regulation, signal transduction, and cell cycle pathways, forming distinct clusters. THCV affected fewer and more dispersed pathways, reflecting narrower molecular engagement. Color scale indicates adjusted p-values.

In contrast, THCV activated a narrower set of pathways, with moderate enrichment in transcriptional activity, cell cycle regulation, and mitotic progression. While THCV did engage signal transduction pathways, the overall intensity and breadth of activation were reduced compared to CBV. This more restricted pattern suggests a selective and possibly short-acting mechanism of action, which aligns with the shorter duration of antinociceptive effects observed in behavioral assays (Ji et al., 2016). Given that THCV acts as a partial agonist or antagonist at cannabinoid receptors depending on dose and receptor context (Pertwee, 2008), its limited transcriptomic footprint may reflect a mechanism primarily mediated through acute receptor interactions rather than broad downstream genomic effects. Notably, both compounds induced minimal pathway suppression, with only minor effects observed in drug ADME and cell cycle categories, suggesting that their impact is largely driven by pathway activation rather than inhibition (Zhao and Iyengar, 2012, Nada et al., 2024).

These findings align with prior biological process enrichment results and reinforce the notion that CBV induces broader and more sustained molecular changes. The strong activation of transcriptional and developmental signaling pathways by CBV may support long-term modulation of nociceptive circuits through mechanisms involving neuroplasticity, chromatin remodeling, and altered synaptic signaling (Denk et al., 2014). Such molecular changes are associated with persistent alterations in pain sensitivity and are often observed in chronic pain models (Zhang et al., 2011, Sun et al., 2012, Denk and McMahon, 2012). In contrast, THCV’s more limited transcriptional footprint could underlie its shorter-acting behavioral effects, possibly due to transient receptor-level engagement and weaker influence on long-term cellular processes (Chen et al., 2020). This divergence further supports distinct therapeutic profiles for each compound and suggests that CBV may be better suited for conditions requiring sustained analgesic action, while THCV may be more appropriate for transient modulation of nociceptive signaling. Taken all together, these pathway-level differences reflect not only distinct pharmacodynamic profiles but also divergent modes of molecular engagement between CBV and THCV. CBV appears to trigger a complex regulatory cascade involving gene expression, developmental signaling, and cellular remodeling, which may translate into prolonged neuromodulatory effects. In contrast, THCV exerts a narrower, possibly more targeted influence, which might offer advantages in clinical contexts where short-term or receptor-specific modulation is desirable. These insights underscore the therapeutic potential of minor cannabinoids and highlight the utility of *C. elegans* as a model for dissecting conserved pathways involved in nociception and cannabinoid pharmacology (Oakes et al., 2019).

Together, our findings demonstrate that CBV and THCV elicit significant dose-dependent antinociceptive effects in *C. elegans*, mediated via distinct interactions with cannabinoid and vanilloid receptors. Complementary proteomic analyses revealed that these compounds modulate different biological processes and signaling pathways. Future studies in mammalian models are needed to validate these results and better characterize the downstream mechanism involved.

## Acknowledgements

The Université de Montréal partially provided financial support for tuition fees to N. Rahmani.

## Supplementary information

A supplementary data file is provided to include experimental schematics and other derived bioinformatic data

## Author’s contributions

NR, JDC, and FB conceived and designed research. NR and JDC conducted experiments and analyzed data. NR, JDC and FB wrote the manuscript. All Authors read, reviewed and approved the manuscript.

## Data availability

The data that support the findings from this study are provided in supplementary files or available from the corresponding author upon reasonable request.

