## Supplemental data for "Cannabivarin and Tetrahydrocannabivarin Modulate Nociception via Vanilloid Channels and Cannabinoid-Like Receptors in *Caenorhabditis elegans*"

**Supplementary Data**

**Figure S1.**  A schematic of the quadrants assay adapted from Margie *et al*. (2013). For head avoidance assay, plates were divided into quadrants two test (A and D) and two controls (B and C). Sodium azide was added to all four quadrants to paralyze nematodes. *C. elegans* were added at the center of the plate (typically, n = 100 to 1,000) and after 30 minutes, animals were counted on each quadrant. Only animals outside the inner circle were scored. The calculation of thermal avoidance index was performed has described.

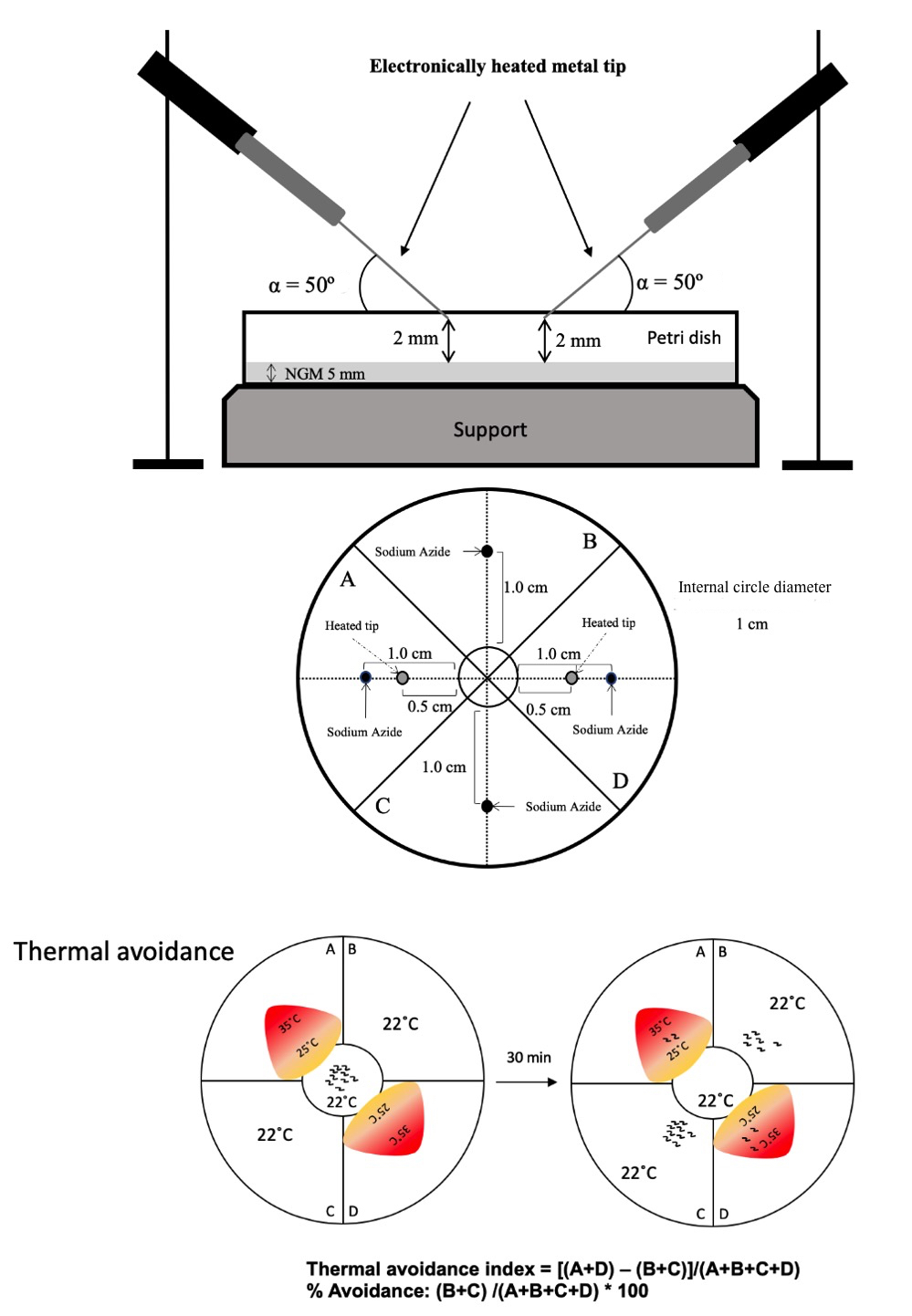

**Table 1. Protein detected and quantified using FragPipe (version 22.0) with the MSFragger search engine for each experimental group.** *C. elegans* reference proteome, obtained from UniProt (taxon identifier 6239) in FASTA format.

| **Protein ID** | **CBV/CTRL log2 FC** | **CBV/CTRL p.val** | **CBV/CTRL p.adj** | **THCV/CTRL log2 FC** | **THCV/CTRL p.val** | **THCV/CTRL p.adj** |
| --- | --- | --- | --- | --- | --- | --- |
| G5EEH9 | 2.85 | 0.0014 | 0.0656 | 3.59 | 0.000334 | 0.0434 |
| G5EGN2 | -2.68 | 0.000517 | 0.0353 | -2.7 | 0.000494 | 0.0473 |
| P52715 | -4.83 | 2.90E-05 | 0.0117 | -4.04 | 9.90E-05 | 0.0286 |
| P55155 | -2.09 | 8.33E-05 | 0.0169 | -1.9 | 0.000156 | 0.0286 |
| P90916 | -0.317 | 0.18 | 0.464 | -1.31 | 0.000357 | 0.0434 |
| Q09531 | 2.52 | 3.86E-05 | 0.0117 | 1.4 | 0.00176 | 0.0731 |
| Q09629 | 0.887 | 0.00717 | 0.121 | 1.37 | 0.000583 | 0.0473 |
| Q10051 | 1.41 | 0.00915 | 0.128 | 2.34 | 0.00053 | 0.0473 |
| Q10663 | 1.49 | 0.000336 | 0.0273 | 0.338 | 0.203 | 0.465 |
| Q17334 | 1.55 | 0.000108 | 0.0169 | 0.863 | 0.00397 | 0.0992 |
| Q9N4X8 | -2.59 | 0.000168 | 0.0204 | -1.33 | 0.00878 | 0.134 |
| B7WN95 | -5.47 | 1.62E-06 | 0.00197 | -5.11 | 2.68E-06 | 0.00326 |
| G5EBP5 | 0.723 | 0.0126 | 0.151 | 1.28 | 0.000525 | 0.0473 |
| G5EDV0 | -2.29 | 0.000111 | 0.0169 | -1.52 | 0.00152 | 0.0731 |
| G5EE67 | 4.08 | 0.000128 | 0.0173 | 0.773 | 0.216 | 0.477 |
| G5EF87 | 3.51 | 8.83E-05 | 0.0169 | 3.53 | 8.52E-05 | 0.0286 |
| G5EFK4 | -0.822 | 0.0341 | 0.217 | -2.19 | 0.000164 | 0.0286 |
| K8ESE2 | -1.96 | 0.000257 | 0.0251 | -2.13 | 0.000148 | 0.0286 |
| O16462 | -2.47 | 2.00E-04 | 0.0221 | -2.26 | 0.000349 | 0.0434 |
| Q18529 | -2.86 | 0.000464 | 0.0353 | -2.39 | 0.00139 | 0.0731 |
| Q19661 | -3.92 | 9.33E-06 | 0.00568 | 0.0796 | 0.838 | 0.922 |
| Q22336 | -2.27 | 0.000522 | 0.0353 | -1.78 | 0.00226 | 0.0758 |
| Q22922 | -3.22 | 0.0036 | 0.0995 | -4.42 | 0.000552 | 0.0473 |
| Q8WSW2 | -2.46 | 0.000323 | 0.0273 | 0.268 | 0.521 | 0.744 |
| Q9BKU3 | 3.64 | 0.000268 | 0.0251 | 4.13 | 0.000115 | 0.0286 |
| A0A0K3AUJ9 | -0.363 | 0.0703 | 0.299 | 0.0391 | 0.826 | 0.917 |
| B1V8A0 | 0.451 | 0.846 | 0.928 | 5.11 | 0.0537 | 0.276 |
| C0HLB3 | 0.671 | 0.021 | 0.189 | 0.21 | 0.391 | 0.653 |
| D0PV95 | 1.02 | 0.0644 | 0.292 | 0.748 | 0.154 | 0.407 |
| G4SLH0 | 0.152 | 0.671 | 0.839 | 0.196 | 0.585 | 0.791 |
| G5ECG0 | -0.332 | 0.597 | 0.798 | 0.299 | 0.635 | 0.821 |
| G5ECM9 | 0.315 | 0.326 | 0.617 | 0.443 | 0.18 | 0.436 |
| G5ECU1 | 2.92 | 0.0416 | 0.23 | 3.22 | 0.0281 | 0.217 |
| G5ECY0 | 0.0869 | 0.719 | 0.867 | 0.418 | 0.112 | 0.358 |
| G5ED41 | 1.19 | 0.0459 | 0.239 | 0.251 | 0.629 | 0.818 |
| G5EDM4 | 0.577 | 0.116 | 0.375 | 1.34 | 0.00374 | 0.0991 |
| G5EDP2 | 0.0301 | 0.95 | 0.973 | 0.264 | 0.584 | 0.791 |
| G5EEE5 | -2.54 | 0.081 | 0.32 | -1.34 | 0.322 | 0.598 |
| G5EEI4 | 0.0416 | 0.889 | 0.941 | -0.296 | 0.337 | 0.612 |
| G5EEK9 | 0.264 | 0.502 | 0.729 | -0.286 | 0.469 | 0.711 |
| G5EFF7 | -0.257 | 0.281 | 0.584 | -0.187 | 0.424 | 0.68 |
| G5EFZ1 | 0.429 | 0.427 | 0.68 | 0.0603 | 0.909 | 0.955 |
| G5EGI7 | 0.488 | 0.398 | 0.661 | -0.128 | 0.82 | 0.914 |
| G5EGK8 | 0.177 | 0.788 | 0.905 | -0.146 | 0.825 | 0.917 |
| G5EGP8 | -0.63 | 0.508 | 0.734 | -0.0433 | 0.963 | 0.986 |
| H2FLJ1 | -1.22 | 0.147 | 0.422 | 0.398 | 0.613 | 0.807 |
| H2KYQ5 | -0.113 | 0.658 | 0.831 | -0.531 | 0.0634 | 0.294 |
| O01504 | -0.649 | 0.0584 | 0.278 | 0.142 | 0.641 | 0.827 |
| O01530 | 0.325 | 0.116 | 0.375 | 0.282 | 0.164 | 0.416 |
| O01541 | 0.194 | 0.812 | 0.911 | -1.71 | 0.0636 | 0.294 |
| O01592 | 2.13 | 0.0423 | 0.23 | 1.73 | 0.0859 | 0.321 |
| O01615 | 1.65 | 0.167 | 0.447 | -0.631 | 0.576 | 0.785 |
| O01692 | -0.167 | 0.43 | 0.682 | -0.395 | 0.0855 | 0.32 |
| O01761 | 0.807 | 0.558 | 0.773 | -0.288 | 0.832 | 0.919 |
| O01789 | -0.0562 | 0.944 | 0.972 | -0.872 | 0.296 | 0.575 |
| O01802 | 0.166 | 0.502 | 0.729 | -0.192 | 0.442 | 0.693 |
| O01803 | -0.321 | 0.722 | 0.868 | -0.597 | 0.513 | 0.742 |
| O01805 | -0.525 | 0.142 | 0.415 | -0.776 | 0.0433 | 0.249 |
| O01812 | -0.434 | 0.129 | 0.395 | -0.162 | 0.544 | 0.761 |
| O01868 | -0.0349 | 0.875 | 0.94 | -0.0182 | 0.935 | 0.972 |
| O01974 | -1.78 | 0.156 | 0.434 | -0.044 | 0.97 | 0.986 |
| O02056 | 0.154 | 0.403 | 0.663 | 0.0303 | 0.866 | 0.933 |
| O02108 | 0.0791 | 0.913 | 0.956 | 1.06 | 0.167 | 0.42 |
| O02115 | 0.371 | 0.363 | 0.646 | 0.496 | 0.234 | 0.497 |
| O02328 | 0.239 | 0.77 | 0.894 | 0.446 | 0.587 | 0.791 |
| O02365 | -0.0333 | 0.88 | 0.94 | 0.0448 | 0.839 | 0.922 |
| O02485 | 1.03 | 0.286 | 0.587 | -0.0191 | 0.984 | 0.991 |
| O02495 | 1.58 | 0.118 | 0.378 | 1.6 | 0.115 | 0.36 |
| O02639 | 0.355 | 0.101 | 0.349 | 0.435 | 0.0526 | 0.271 |
| O02640 | 0.0464 | 0.787 | 0.905 | 0.00936 | 0.956 | 0.985 |
| O16202 | 0.791 | 0.394 | 0.661 | 0.982 | 0.296 | 0.575 |
| O16259 | 2.2 | 0.0141 | 0.159 | 2.78 | 0.00432 | 0.0992 |
| O16264 | -0.58 | 0.0688 | 0.298 | -0.566 | 0.0748 | 0.311 |
| O16294 | 1.76 | 0.219 | 0.516 | 2.46 | 0.0998 | 0.337 |
| O16305 | -0.399 | 0.398 | 0.661 | 0.246 | 0.597 | 0.798 |
| O16368 | 1.3 | 0.269 | 0.571 | 0.741 | 0.515 | 0.742 |
| O17071 | 1.12 | 0.176 | 0.458 | 0.856 | 0.289 | 0.565 |
| O17214 | -0.0808 | 0.666 | 0.836 | 0.054 | 0.773 | 0.89 |
| O17271 | -0.672 | 0.61 | 0.808 | 0.25 | 0.849 | 0.926 |
| O17389 | 0.127 | 0.624 | 0.813 | -0.0631 | 0.806 | 0.904 |
| O17536 | 0.306 | 0.825 | 0.917 | -0.179 | 0.897 | 0.949 |
| O17570 | 0.3 | 0.387 | 0.66 | 0.61 | 0.101 | 0.339 |
| O17586 | -0.278 | 0.492 | 0.721 | -0.154 | 0.701 | 0.854 |
| O17607 | 2.17 | 0.143 | 0.417 | 2.9 | 0.0632 | 0.294 |
| O17622 | -1.9 | 0.144 | 0.417 | -1.64 | 0.198 | 0.458 |
| O17680 | -0.151 | 0.402 | 0.662 | -0.357 | 0.0713 | 0.305 |
| O17695 | 1.89 | 0.055 | 0.27 | 0.49 | 0.575 | 0.785 |
| O17732 | 1.64 | 0.193 | 0.481 | -0.179 | 0.88 | 0.942 |
| O17861 | -0.048 | 0.82 | 0.916 | 0.0963 | 0.65 | 0.831 |
| O17915 | 0.232 | 0.351 | 0.637 | -0.233 | 0.348 | 0.622 |
| O17919 | 0.622 | 0.0324 | 0.214 | 0.285 | 0.268 | 0.541 |
| O17953 | -0.185 | 0.415 | 0.671 | 0.208 | 0.363 | 0.633 |
| O18054 | 0.78 | 0.491 | 0.721 | -0.0339 | 0.976 | 0.986 |
| O18650 | 0.141 | 0.453 | 0.696 | 0.114 | 0.54 | 0.757 |
| O44156 | 0.744 | 0.0381 | 0.226 | 0.82 | 0.0259 | 0.216 |
| O44400 | 1.05 | 0.00829 | 0.123 | 0.507 | 0.127 | 0.372 |
| O44411 | 1.08 | 0.479 | 0.711 | 1.59 | 0.308 | 0.588 |
| O44441 | -0.454 | 0.0627 | 0.29 | -0.21 | 0.345 | 0.621 |
| O44443 | 0.564 | 0.567 | 0.775 | -0.0634 | 0.948 | 0.983 |
| O44451 | -0.536 | 0.643 | 0.821 | -0.363 | 0.753 | 0.882 |
| O44480 | 0.0524 | 0.76 | 0.888 | -0.0845 | 0.624 | 0.813 |
| O44572 | 1.26 | 0.168 | 0.448 | -1.43 | 0.126 | 0.372 |
| O44739 | -0.597 | 0.392 | 0.661 | -1.56 | 0.0465 | 0.257 |
| O45181 | 0.551 | 0.0666 | 0.294 | 0.0455 | 0.864 | 0.933 |
| O45319 | -1.7 | 0.00272 | 0.0841 | -0.997 | 0.035 | 0.232 |
| O45346 | -2.47 | 0.0155 | 0.165 | 0.102 | 0.902 | 0.95 |
| O45495 | -0.518 | 0.305 | 0.604 | 0.227 | 0.644 | 0.828 |
| O45499 | 0.19 | 0.343 | 0.636 | 0.289 | 0.164 | 0.416 |
| O45679 | -0.598 | 0.0227 | 0.196 | -0.755 | 0.00762 | 0.131 |
| O45734 | -0.241 | 0.336 | 0.632 | -0.0131 | 0.957 | 0.985 |
| O45924 | 0.15 | 0.801 | 0.909 | -0.477 | 0.43 | 0.686 |
| O45946 | -0.096 | 0.627 | 0.813 | -0.113 | 0.569 | 0.782 |
| O61199 | 0.0878 | 0.769 | 0.893 | -0.267 | 0.383 | 0.65 |
| O61235 | 1.5 | 0.122 | 0.383 | 0.229 | 0.797 | 0.902 |
| O61708 | -0.558 | 0.0763 | 0.313 | -0.673 | 0.0401 | 0.238 |
| O61955 | 0.354 | 0.761 | 0.888 | 2.33 | 0.0735 | 0.309 |
| O62053 | -0.234 | 0.541 | 0.761 | 0.256 | 0.503 | 0.737 |
| O62106 | 0.559 | 0.412 | 0.67 | 1.86 | 0.0214 | 0.199 |
| O62140 | -0.305 | 0.626 | 0.813 | 0.704 | 0.277 | 0.551 |
| O62146 | -0.862 | 0.0214 | 0.19 | -0.403 | 0.216 | 0.477 |
| O62220 | 0.166 | 0.51 | 0.734 | 0.201 | 0.429 | 0.685 |
| O62246 | -2.11 | 0.0815 | 0.32 | 0.403 | 0.713 | 0.859 |
| O62327 | 0.0922 | 0.673 | 0.84 | -0.0946 | 0.665 | 0.838 |
| O76360 | 1.3 | 0.0645 | 0.292 | -0.266 | 0.672 | 0.839 |
| O76371 | 0.0717 | 0.823 | 0.916 | 0.278 | 0.397 | 0.657 |
| O76577 | -0.299 | 0.515 | 0.737 | -1.01 | 0.0519 | 0.27 |
| O76840 | 0.0378 | 0.874 | 0.939 | 0.272 | 0.272 | 0.545 |
| P02566 | 0.156 | 0.435 | 0.683 | -0.0819 | 0.677 | 0.841 |
| P02567 | 0.898 | 0.00211 | 0.0825 | 0.154 | 0.457 | 0.703 |
| P03934 | 0.0523 | 0.822 | 0.916 | -0.0388 | 0.867 | 0.933 |
| P04970 | -0.206 | 0.524 | 0.743 | -0.23 | 0.477 | 0.712 |
| P05634 | -1 | 0.00907 | 0.128 | -0.585 | 0.0786 | 0.312 |
| P05690 | -0.645 | 0.0366 | 0.221 | -0.808 | 0.0141 | 0.16 |
| P06125 | -1.06 | 0.0424 | 0.23 | -1.18 | 0.0281 | 0.217 |
| P08898 | -0.0684 | 0.814 | 0.912 | -0.233 | 0.433 | 0.686 |
| P09446 | -0.618 | 0.0129 | 0.151 | -0.42 | 0.061 | 0.292 |
| P09588 | 0.82 | 0.0257 | 0.201 | -0.086 | 0.779 | 0.895 |
| P0DM41 | -0.577 | 0.614 | 0.809 | 2.44 | 0.0588 | 0.289 |
| P10299 | 0.0406 | 0.883 | 0.941 | -0.104 | 0.708 | 0.859 |
| P10567 | -0.0869 | 0.628 | 0.813 | 0.146 | 0.421 | 0.679 |
| P10771 | 0.511 | 0.0383 | 0.226 | 0.269 | 0.227 | 0.489 |
| P11141 | -0.223 | 0.315 | 0.612 | 0.122 | 0.573 | 0.783 |
| P12844 | 0.349 | 0.134 | 0.401 | 0.235 | 0.293 | 0.571 |
| P12845 | 0.656 | 0.015 | 0.163 | -0.018 | 0.934 | 0.972 |
| P15796 | 0.878 | 0.167 | 0.447 | -0.148 | 0.803 | 0.902 |
| P17140 | 0.209 | 0.562 | 0.774 | 0.551 | 0.15 | 0.402 |
| P17329 | -0.725 | 0.158 | 0.434 | -1.87 | 0.00413 | 0.0992 |
| P17330 | 0.167 | 0.384 | 0.659 | 0.186 | 0.334 | 0.61 |
| P17343 | 0.79 | 0.316 | 0.612 | 1.13 | 0.164 | 0.416 |
| P17512 | 3.23 | 0.0556 | 0.271 | 0.575 | 0.699 | 0.854 |
| P18334 | 1.73 | 0.00484 | 0.107 | 2.24 | 0.00109 | 0.0731 |
| P18947 | -1.1 | 0.00462 | 0.107 | -0.752 | 0.028 | 0.217 |
| P18948 | -0.313 | 0.227 | 0.527 | -0.472 | 0.0841 | 0.318 |
| P19625 | -0.29 | 0.18 | 0.464 | -0.13 | 0.528 | 0.747 |
| P19826 | 0.683 | 0.325 | 0.617 | -0.389 | 0.566 | 0.78 |
| P19974 | -0.463 | 0.0328 | 0.214 | -0.378 | 0.0682 | 0.301 |
| P20163 | -0.31 | 0.409 | 0.67 | -0.385 | 0.312 | 0.59 |
| P24894 | -0.0127 | 0.958 | 0.978 | -0.094 | 0.698 | 0.854 |
| P25807 | 1.94 | 0.0839 | 0.325 | 1.03 | 0.322 | 0.598 |
| P27420 | -0.443 | 0.0311 | 0.212 | -0.148 | 0.404 | 0.662 |
| P27604 | -0.396 | 0.176 | 0.459 | -0.103 | 0.709 | 0.859 |
| P27639 | -0.22 | 0.427 | 0.681 | -0.2 | 0.47 | 0.711 |
| P27798 | -0.559 | 0.011 | 0.139 | -0.169 | 0.344 | 0.62 |
| P29691 | -0.126 | 0.6 | 0.799 | -0.463 | 0.0813 | 0.312 |
| P30625 | 0.184 | 0.645 | 0.821 | 0.213 | 0.595 | 0.798 |
| P30627 | -1.36 | 0.00572 | 0.109 | -1.59 | 0.00249 | 0.0777 |
| P30629 | 0.101 | 0.724 | 0.869 | 0.697 | 0.0372 | 0.235 |
| P30632 | 0.964 | 0.0631 | 0.291 | 0.339 | 0.468 | 0.711 |
| P30642 | 0.496 | 0.151 | 0.428 | -0.725 | 0.0494 | 0.265 |
| P31161 | -0.188 | 0.697 | 0.852 | -0.237 | 0.624 | 0.813 |
| P34255 | 0.818 | 0.0397 | 0.228 | 0.451 | 0.211 | 0.473 |
| P34286 | -0.454 | 0.679 | 0.843 | -0.361 | 0.741 | 0.879 |
| P34328 | -0.522 | 0.086 | 0.331 | -0.474 | 0.113 | 0.358 |
| P34329 | 0.311 | 0.405 | 0.666 | 0.555 | 0.156 | 0.409 |
| P34334 | 0.214 | 0.361 | 0.645 | 0.0654 | 0.775 | 0.891 |
| P34339 | -1.45 | 0.0695 | 0.299 | -0.57 | 0.431 | 0.686 |
| P34346 | 0.293 | 0.35 | 0.637 | 0.522 | 0.117 | 0.362 |
| P34382 | -0.386 | 0.143 | 0.417 | -0.143 | 0.562 | 0.778 |
| P34383 | -0.604 | 0.0789 | 0.314 | -0.236 | 0.453 | 0.703 |
| P34385 | -1.52 | 0.0131 | 0.151 | -1.15 | 0.0415 | 0.244 |
| P34445 | 0.848 | 0.413 | 0.67 | 0.882 | 0.395 | 0.655 |
| P34446 | 2.06 | 0.0645 | 0.292 | 1.94 | 0.0778 | 0.311 |
| P34447 | -0.0442 | 0.946 | 0.972 | -0.676 | 0.314 | 0.592 |
| P34455 | 0.573 | 0.0262 | 0.203 | 0.181 | 0.411 | 0.669 |
| P34460 | 0.0108 | 0.965 | 0.982 | 0.309 | 0.235 | 0.497 |
| P34462 | 0.378 | 0.169 | 0.449 | 0.371 | 0.176 | 0.432 |
| P34466 | -0.651 | 0.238 | 0.538 | -0.857 | 0.132 | 0.382 |
| P34496 | 0.29 | 0.482 | 0.713 | 0.765 | 0.0889 | 0.327 |
| P34500 | -0.0956 | 0.829 | 0.92 | -0.147 | 0.741 | 0.879 |
| P34517 | -0.421 | 0.603 | 0.802 | -0.755 | 0.361 | 0.632 |
| P34519 | 1.33 | 0.302 | 0.603 | 1.31 | 0.308 | 0.588 |
| P34526 | -1.17 | 0.0993 | 0.349 | -1.16 | 0.102 | 0.339 |
| P34528 | -0.677 | 0.346 | 0.636 | 0.86 | 0.239 | 0.503 |
| P34539 | 0.787 | 0.263 | 0.564 | -0.359 | 0.597 | 0.798 |
| P34540 | -0.895 | 0.644 | 0.821 | -2.65 | 0.195 | 0.455 |
| P34559 | -1.21 | 0.24 | 0.538 | 0.00962 | 0.992 | 0.997 |
| P34563 | 0.384 | 0.336 | 0.632 | 1.08 | 0.0216 | 0.199 |
| P34574 | 1.16 | 0.00415 | 0.103 | 0.545 | 0.0957 | 0.335 |
| P34575 | -0.0599 | 0.863 | 0.936 | -0.539 | 0.149 | 0.401 |
| P34594 | 0.317 | 0.545 | 0.762 | 0.265 | 0.611 | 0.806 |
| P34618 | 0.216 | 0.881 | 0.941 | -1.77 | 0.243 | 0.504 |
| P34629 | -0.00237 | 0.997 | 0.998 | -0.501 | 0.468 | 0.711 |
| P34645 | -2.96 | 0.0128 | 0.151 | -1.43 | 0.159 | 0.41 |
| P34654 | -2.42 | 0.116 | 0.375 | -2.27 | 0.136 | 0.385 |
| P34662 | 0.0941 | 0.667 | 0.837 | 0.311 | 0.18 | 0.436 |
| P34685 | 0.617 | 0.0789 | 0.314 | 0.864 | 0.023 | 0.202 |
| P34689 | 0.698 | 0.591 | 0.792 | -1.52 | 0.258 | 0.522 |
| P34690 | -1.83 | 0.0431 | 0.231 | -1.07 | 0.196 | 0.456 |
| P34697 | -0.67 | 0.0222 | 0.195 | -0.0435 | 0.857 | 0.931 |
| P34714 | -1.14 | 0.0272 | 0.208 | -0.641 | 0.166 | 0.418 |
| P35129 | -1.47 | 0.147 | 0.423 | -0.9 | 0.354 | 0.626 |
| P36573 | 0.0629 | 0.826 | 0.917 | -0.00897 | 0.975 | 0.986 |
| P36609 | -0.829 | 0.339 | 0.632 | 0.376 | 0.657 | 0.835 |
| P37165 | -0.272 | 0.387 | 0.66 | -0.271 | 0.389 | 0.653 |
| P37806 | -0.429 | 0.214 | 0.514 | -0.716 | 0.0555 | 0.277 |
| P39055 | -3.66 | 0.0398 | 0.228 | -3.93 | 0.0299 | 0.224 |
| P40614 | 0.00608 | 0.985 | 0.994 | -0.247 | 0.46 | 0.705 |
| P41847 | -0.337 | 0.435 | 0.683 | -0.656 | 0.149 | 0.401 |
| P41932 | 0.276 | 0.175 | 0.458 | 0.868 | 0.00176 | 0.0731 |
| P41938 | 0.387 | 0.0976 | 0.349 | 0.43 | 0.0707 | 0.303 |
| P41988 | -0.18 | 0.702 | 0.853 | -0.342 | 0.471 | 0.712 |
| P41994 | 0.859 | 0.705 | 0.855 | 0.233 | 0.918 | 0.963 |
| P41996 | 0.46 | 0.458 | 0.697 | -0.182 | 0.765 | 0.887 |
| P43508 | 2.13 | 0.00666 | 0.117 | 2.33 | 0.00418 | 0.0992 |
| P43510 | 0.236 | 0.563 | 0.774 | 0.703 | 0.111 | 0.357 |
| P45971 | 1.09 | 0.271 | 0.573 | 0.355 | 0.711 | 0.859 |
| P46502 | 0.294 | 0.355 | 0.638 | 0.353 | 0.272 | 0.545 |
| P46548 | 0.2 | 0.804 | 0.91 | 1.07 | 0.208 | 0.47 |
| P46550 | 0.948 | 0.0919 | 0.34 | 0.338 | 0.512 | 0.742 |
| P46561 | 0.179 | 0.437 | 0.684 | -0.405 | 0.102 | 0.339 |
| P46562 | 0.209 | 0.464 | 0.699 | -0.364 | 0.219 | 0.48 |
| P46563 | 0.526 | 0.0216 | 0.191 | 0.319 | 0.121 | 0.367 |
| P46769 | 0.114 | 0.646 | 0.822 | 0.116 | 0.641 | 0.827 |
| P46822 | 0.49 | 0.429 | 0.681 | 0.248 | 0.684 | 0.847 |
| P46975 | 1.8 | 0.0214 | 0.19 | -1.69 | 0.0276 | 0.217 |
| P47207 | 0.481 | 0.121 | 0.381 | 0.512 | 0.102 | 0.339 |
| P47208 | 0.662 | 0.0707 | 0.299 | 0.0432 | 0.895 | 0.949 |
| P47209 | 0.198 | 0.379 | 0.658 | 0.0839 | 0.703 | 0.854 |
| P47991 | 0.0752 | 0.672 | 0.84 | -0.15 | 0.409 | 0.667 |
| P48150 | 0.279 | 0.187 | 0.471 | 0.194 | 0.343 | 0.619 |
| P48152 | 0.116 | 0.575 | 0.78 | 0.0202 | 0.921 | 0.964 |
| P48154 | 0.0593 | 0.728 | 0.869 | -0.161 | 0.359 | 0.631 |
| P48156 | 0.345 | 0.13 | 0.395 | 0.173 | 0.421 | 0.679 |
| P48158 | 0.425 | 0.0354 | 0.221 | 0.0663 | 0.701 | 0.854 |
| P48162 | -0.293 | 0.502 | 0.729 | 0.252 | 0.562 | 0.778 |
| P48166 | 0.0223 | 0.96 | 0.979 | 0.291 | 0.521 | 0.744 |
| P49041 | -0.193 | 0.397 | 0.661 | -0.485 | 0.056 | 0.277 |
| P49180 | -0.135 | 0.54 | 0.761 | 0.0991 | 0.651 | 0.831 |
| P49181 | 0.257 | 0.264 | 0.566 | 0.321 | 0.171 | 0.425 |
| P49196 | -0.463 | 0.0417 | 0.23 | 0.266 | 0.199 | 0.459 |
| P49197 | -0.128 | 0.589 | 0.79 | 0.463 | 0.0778 | 0.311 |
| P49405 | -0.0523 | 0.793 | 0.908 | -0.287 | 0.176 | 0.432 |
| P49595 | 2.85 | 0.00309 | 0.0912 | 1.77 | 0.0307 | 0.224 |
| P49632 | -0.175 | 0.54 | 0.761 | 0.0892 | 0.753 | 0.882 |
| P50093 | 0.4 | 0.1 | 0.349 | 0.19 | 0.402 | 0.661 |
| P50140 | 0.158 | 0.393 | 0.661 | 0.3 | 0.125 | 0.372 |
| P50432 | -0.192 | 0.368 | 0.651 | -0.369 | 0.106 | 0.347 |
| P50880 | 0.128 | 0.449 | 0.695 | 0.155 | 0.364 | 0.635 |
| P51403 | 0.107 | 0.544 | 0.762 | -0.343 | 0.0769 | 0.311 |
| P51404 | 0.33 | 0.116 | 0.375 | 0.0197 | 0.918 | 0.963 |
| P51875 | 0.929 | 0.192 | 0.481 | 0.515 | 0.451 | 0.702 |
| P52009 | -0.493 | 0.0342 | 0.217 | -0.181 | 0.373 | 0.642 |
| P52011 | 0.124 | 0.551 | 0.766 | 0.0544 | 0.792 | 0.9 |
| P52013 | -0.47 | 0.067 | 0.294 | -0.333 | 0.171 | 0.425 |
| P52014 | -1.24 | 0.111 | 0.367 | 0.239 | 0.738 | 0.878 |
| P52015 | 0.277 | 0.281 | 0.584 | 0.161 | 0.521 | 0.744 |
| P52057 | -3.31 | 0.0599 | 0.283 | -3.15 | 0.0702 | 0.302 |
| P52275 | -0.479 | 0.107 | 0.361 | -0.961 | 0.00697 | 0.127 |
| P52554 | -2.35 | 0.0998 | 0.349 | -2.67 | 0.0681 | 0.301 |
| P52709 | -0.228 | 0.526 | 0.744 | -0.228 | 0.526 | 0.746 |
| P52713 | 0.421 | 0.0539 | 0.268 | 0.157 | 0.42 | 0.679 |
| P52716 | -0.873 | 0.464 | 0.699 | -0.519 | 0.659 | 0.837 |
| P52717 | -0.142 | 0.575 | 0.78 | 0.0158 | 0.95 | 0.983 |
| P52814 | -0.263 | 0.392 | 0.661 | -0.561 | 0.0914 | 0.33 |
| P52819 | -0.393 | 0.104 | 0.357 | -0.47 | 0.06 | 0.291 |
| P52821 | 0.0784 | 0.908 | 0.951 | 0.248 | 0.714 | 0.859 |
| P52899 | 0.892 | 0.0967 | 0.348 | 0.867 | 0.105 | 0.344 |
| P53013 | -0.161 | 0.382 | 0.659 | -0.22 | 0.243 | 0.504 |
| P53014 | -0.542 | 0.0905 | 0.336 | -0.29 | 0.332 | 0.609 |
| P53017 | 0.00926 | 0.991 | 0.998 | 1.63 | 0.07 | 0.302 |
| P53585 | 0.587 | 0.148 | 0.423 | -0.474 | 0.231 | 0.494 |
| P53588 | -0.0281 | 0.971 | 0.985 | -1.92 | 0.0352 | 0.232 |
| P53589 | 1.35 | 0.24 | 0.538 | -0.797 | 0.473 | 0.712 |
| P53596 | 0.193 | 0.303 | 0.603 | 0.0509 | 0.779 | 0.895 |
| P53703 | 1.02 | 0.325 | 0.617 | 0.8 | 0.436 | 0.687 |
| P54216 | 0.0594 | 0.835 | 0.923 | 0.0172 | 0.952 | 0.983 |
| P54412 | 0.18 | 0.421 | 0.676 | 0.00181 | 0.993 | 0.997 |
| P54688 | -1.01 | 0.239 | 0.538 | -1.87 | 0.0479 | 0.26 |
| P54811 | 0.736 | 0.0108 | 0.138 | 0.0723 | 0.75 | 0.882 |
| P54812 | 1.19 | 0.111 | 0.367 | -0.12 | 0.861 | 0.931 |
| P54889 | 0.09 | 0.905 | 0.95 | 1.27 | 0.123 | 0.372 |
| P55216 | -2.63 | 0.112 | 0.368 | -0.407 | 0.789 | 0.898 |
| P55326 | -0.0801 | 0.811 | 0.911 | -0.0653 | 0.846 | 0.925 |
| P55853 | -1.38 | 0.0252 | 0.2 | -0.997 | 0.0805 | 0.312 |
| P55954 | -0.32 | 0.279 | 0.583 | -0.361 | 0.227 | 0.489 |
| P55955 | 1.09 | 0.00807 | 0.121 | 0.924 | 0.0179 | 0.186 |
| P55956 | 2.05 | 0.247 | 0.545 | 1.62 | 0.353 | 0.625 |
| P61866 | -0.0599 | 0.731 | 0.869 | -0.0532 | 0.76 | 0.886 |
| P62784 | -0.189 | 0.603 | 0.802 | -0.544 | 0.158 | 0.41 |
| P83351 | 1.38 | 0.071 | 0.299 | 1.41 | 0.0651 | 0.297 |
| P90795 | 1.52 | 0.105 | 0.358 | 0.805 | 0.358 | 0.631 |
| P90850 | 0.618 | 0.343 | 0.636 | 0.576 | 0.375 | 0.643 |
| P90897 | 0.00351 | 0.994 | 0.998 | -0.576 | 0.227 | 0.489 |
| P90900 | 0.68 | 0.31 | 0.609 | 0.29 | 0.656 | 0.835 |
| P90921 | 0.158 | 0.771 | 0.894 | -0.766 | 0.184 | 0.443 |
| P90978 | -0.0338 | 0.978 | 0.989 | -2.17 | 0.105 | 0.344 |
| P90994 | -0.309 | 0.838 | 0.924 | -0.377 | 0.802 | 0.902 |
| P91027 | 0.0164 | 0.934 | 0.966 | -0.134 | 0.505 | 0.737 |
| P91127 | -0.364 | 0.413 | 0.67 | -0.189 | 0.666 | 0.838 |
| P91128 | 0.111 | 0.562 | 0.774 | 0.115 | 0.549 | 0.766 |
| P91134 | 0.155 | 0.748 | 0.878 | -0.46 | 0.354 | 0.626 |
| P91253 | -0.331 | 0.283 | 0.585 | -1.05 | 0.00707 | 0.127 |
| P91277 | -0.219 | 0.705 | 0.855 | -0.45 | 0.444 | 0.694 |
| P91303 | 0.368 | 0.174 | 0.458 | 0.729 | 0.0189 | 0.189 |
| P91374 | 0.261 | 0.239 | 0.538 | 0.0582 | 0.783 | 0.896 |
| P91427 | 0.51 | 0.0696 | 0.299 | 0.161 | 0.526 | 0.746 |
| P91477 | 0.26 | 0.552 | 0.766 | -0.706 | 0.131 | 0.38 |
| P91859 | 0.575 | 0.124 | 0.387 | 0.881 | 0.0306 | 0.224 |
| P91913 | -0.401 | 0.231 | 0.533 | 0.517 | 0.134 | 0.384 |
| P91914 | 0.313 | 0.299 | 0.601 | -0.0833 | 0.775 | 0.891 |
| P91917 | 0.0543 | 0.856 | 0.935 | -0.764 | 0.0307 | 0.224 |
| P98080 | 0.0553 | 0.752 | 0.88 | 0.174 | 0.334 | 0.61 |
| Q03565 | 0.707 | 0.384 | 0.659 | 0.765 | 0.349 | 0.622 |
| Q03577 | 1.75 | 0.0991 | 0.349 | 2.54 | 0.0273 | 0.217 |
| Q04908 | 0.939 | 0.0363 | 0.221 | 0.705 | 0.0953 | 0.334 |
| Q05036 | 0.433 | 0.149 | 0.424 | -0.146 | 0.605 | 0.802 |
| Q07749 | -1.49 | 0.00768 | 0.121 | -2.02 | 0.00147 | 0.0731 |
| Q07750 | 1.32 | 0.0185 | 0.184 | 0.555 | 0.246 | 0.508 |
| Q09165 | 0.206 | 0.68 | 0.843 | 0.678 | 0.198 | 0.458 |
| Q09227 | -0.768 | 0.389 | 0.661 | 0.147 | 0.866 | 0.933 |
| Q09237 | -0.0775 | 0.809 | 0.911 | 0.409 | 0.225 | 0.487 |
| Q09248 | 0.312 | 0.306 | 0.606 | -1.39 | 0.00138 | 0.0731 |
| Q09250 | 0.401 | 0.55 | 0.765 | 0.391 | 0.56 | 0.777 |
| Q09254 | 2.32 | 0.122 | 0.383 | 3.03 | 0.0546 | 0.277 |
| Q09289 | 0.221 | 0.411 | 0.67 | 0.54 | 0.0682 | 0.301 |
| Q09359 | 0.51 | 0.584 | 0.787 | -0.823 | 0.384 | 0.65 |
| Q09364 | 1.21 | 0.112 | 0.368 | 3.39 | 0.00121 | 0.0731 |
| Q09365 | -1.28 | 0.00388 | 0.101 | -0.923 | 0.0196 | 0.189 |
| Q09390 | -1.19 | 0.45 | 0.695 | -1.83 | 0.257 | 0.521 |
| Q09476 | -0.14 | 0.523 | 0.743 | 0.196 | 0.378 | 0.646 |
| Q09508 | 0.612 | 0.0184 | 0.183 | 0.716 | 0.00882 | 0.134 |
| Q09510 | -0.122 | 0.834 | 0.923 | -0.423 | 0.473 | 0.712 |
| Q09511 | 0.0385 | 0.924 | 0.963 | 0.997 | 0.0359 | 0.233 |
| Q09533 | 0.163 | 0.439 | 0.685 | 0.0561 | 0.786 | 0.896 |
| Q09543 | 0.478 | 0.313 | 0.612 | -0.59 | 0.22 | 0.481 |
| Q09544 | -0.783 | 0.00977 | 0.134 | -0.385 | 0.132 | 0.382 |
| Q09545 | 0.503 | 0.0568 | 0.273 | 0.944 | 0.00332 | 0.0917 |
| Q09581 | 0.561 | 0.039 | 0.227 | 0.673 | 0.0185 | 0.188 |
| Q09583 | 0.256 | 0.56 | 0.774 | 0.604 | 0.189 | 0.448 |
| Q09596 | 0.0612 | 0.791 | 0.907 | -0.157 | 0.501 | 0.736 |
| Q09607 | 1.04 | 0.401 | 0.662 | 0.33 | 0.785 | 0.896 |
| Q09610 | 0.497 | 0.0737 | 0.305 | 0.435 | 0.109 | 0.351 |
| Q09665 | -0.476 | 0.125 | 0.387 | -0.296 | 0.316 | 0.594 |
| Q09936 | -0.292 | 0.239 | 0.538 | -0.0472 | 0.842 | 0.924 |
| Q09958 | -1.68 | 0.195 | 0.483 | -2.04 | 0.125 | 0.372 |
| Q0G819 | 1.58 | 0.14 | 0.412 | 1.55 | 0.147 | 0.4 |
| Q10013 | -0.129 | 0.863 | 0.936 | 1.13 | 0.159 | 0.41 |
| Q10020 | 0.675 | 0.0251 | 0.2 | 1.1 | 0.00215 | 0.0758 |
| Q10021 | 0.135 | 0.897 | 0.945 | 0.599 | 0.57 | 0.782 |
| Q10033 | -0.409 | 0.744 | 0.876 | 0.952 | 0.455 | 0.703 |
| Q10121 | -0.262 | 0.359 | 0.645 | 0.654 | 0.0426 | 0.248 |
| Q10454 | 0.151 | 0.374 | 0.654 | 0.0936 | 0.576 | 0.785 |
| Q10576 | -0.416 | 0.481 | 0.712 | 0.665 | 0.272 | 0.545 |
| Q10657 | -0.0325 | 0.891 | 0.942 | -0.366 | 0.152 | 0.406 |
| Q10661 | -0.136 | 0.741 | 0.875 | 0.773 | 0.0898 | 0.328 |
| Q10943 | -1.53 | 0.134 | 0.401 | -0.74 | 0.441 | 0.692 |
| Q11067 | -0.151 | 0.625 | 0.813 | -0.0465 | 0.88 | 0.942 |
| Q11117 | 0.674 | 0.54 | 0.761 | 0.369 | 0.735 | 0.877 |
| Q11176 | -1.79 | 0.041 | 0.229 | -2.44 | 0.0109 | 0.141 |
| Q17335 | 0.529 | 0.437 | 0.684 | -1.06 | 0.143 | 0.395 |
| Q17348 | 0.217 | 0.684 | 0.845 | -0.157 | 0.768 | 0.889 |
| Q17413 | 1.03 | 0.0933 | 0.342 | -0.266 | 0.635 | 0.821 |
| Q17435 | 0.233 | 0.865 | 0.937 | 2.09 | 0.155 | 0.407 |
| Q17460 | -1.99 | 0.00514 | 0.107 | -2.47 | 0.00151 | 0.0731 |
| Q17684 | -0.0624 | 0.932 | 0.966 | 0.322 | 0.66 | 0.837 |
| Q17754 | 0.0693 | 0.87 | 0.939 | -0.0292 | 0.945 | 0.981 |
| Q17770 | -0.534 | 0.0105 | 0.138 | -0.304 | 0.0934 | 0.331 |
| Q17802 | 0.485 | 0.384 | 0.659 | 0.772 | 0.181 | 0.437 |
| Q17827 | -0.323 | 0.325 | 0.617 | -0.2 | 0.534 | 0.751 |
| Q17886 | 2.61 | 0.0145 | 0.16 | 1.23 | 0.177 | 0.433 |
| Q17967 | -0.165 | 0.406 | 0.666 | 0.182 | 0.361 | 0.632 |
| Q18012 | 0.237 | 0.523 | 0.743 | 0.621 | 0.12 | 0.367 |
| Q18026 | -2.05 | 0.00528 | 0.107 | -2.37 | 0.00234 | 0.0758 |
| Q18040 | -0.459 | 0.391 | 0.661 | -0.865 | 0.127 | 0.372 |
| Q18066 | -0.265 | 0.228 | 0.527 | -0.354 | 0.119 | 0.366 |
| Q18090 | -0.465 | 0.415 | 0.671 | -0.8 | 0.178 | 0.435 |
| Q18115 | -1.16 | 0.352 | 0.637 | -0.563 | 0.644 | 0.828 |
| Q18164 | 0.498 | 0.0304 | 0.212 | 0.379 | 0.08 | 0.312 |
| Q18212 | 0.336 | 0.618 | 0.811 | -0.663 | 0.337 | 0.612 |
| Q18240 | -0.721 | 0.165 | 0.445 | -0.326 | 0.508 | 0.741 |
| Q18359 | -0.572 | 0.097 | 0.348 | -0.00916 | 0.977 | 0.986 |
| Q18409 | 0.182 | 0.793 | 0.908 | -0.138 | 0.843 | 0.924 |
| Q18421 | -0.173 | 0.877 | 0.94 | -1.24 | 0.285 | 0.562 |
| Q18678 | 1.14 | 0.0192 | 0.186 | 1.14 | 0.0192 | 0.189 |
| Q18680 | -0.542 | 0.152 | 0.428 | -0.262 | 0.465 | 0.709 |
| Q18688 | 0.226 | 0.342 | 0.636 | 0.0937 | 0.686 | 0.847 |
| Q18785 | -0.969 | 0.000923 | 0.0538 | -0.532 | 0.0217 | 0.199 |
| Q18786 | -1.56 | 0.104 | 0.356 | -0.444 | 0.613 | 0.807 |
| Q18787 | 0.962 | 0.316 | 0.612 | -0.355 | 0.703 | 0.854 |
| Q18803 | 0.475 | 0.0906 | 0.336 | -0.826 | 0.0105 | 0.141 |
| Q18823 | 1.63 | 0.0711 | 0.299 | 1 | 0.233 | 0.497 |
| Q18885 | -0.0532 | 0.783 | 0.903 | 0.0433 | 0.822 | 0.915 |
| Q18938 | -0.508 | 0.42 | 0.676 | -0.334 | 0.592 | 0.794 |
| Q19052 | 1.09 | 0.213 | 0.514 | -0.163 | 0.844 | 0.925 |
| Q19087 | -0.308 | 0.309 | 0.608 | -0.322 | 0.288 | 0.565 |
| Q19126 | -0.0821 | 0.8 | 0.909 | -0.18 | 0.582 | 0.79 |
| Q19162 | 0.149 | 0.433 | 0.683 | 0.0965 | 0.606 | 0.802 |
| Q19191 | -0.259 | 0.579 | 0.784 | -0.743 | 0.137 | 0.386 |
| Q19264 | -1.23 | 0.0363 | 0.221 | -1.45 | 0.0179 | 0.186 |
| Q19286 | 0.806 | 0.0426 | 0.23 | 0.7 | 0.0695 | 0.302 |
| Q19289 | -0.117 | 0.545 | 0.762 | 0.184 | 0.35 | 0.623 |
| Q19375 | -0.476 | 0.0648 | 0.292 | -0.0623 | 0.785 | 0.896 |
| Q19420 | -0.996 | 0.141 | 0.413 | -2.12 | 0.00884 | 0.134 |
| Q19584 | -2.6 | 0.0137 | 0.156 | -2.88 | 0.00845 | 0.134 |
| Q19626 | 0.275 | 0.3 | 0.601 | 0.143 | 0.58 | 0.789 |
| Q19722 | 0.503 | 0.562 | 0.774 | 0.847 | 0.339 | 0.613 |
| Q19724 | -2.24 | 0.118 | 0.378 | -2.48 | 0.0888 | 0.327 |
| Q19743 | 2 | 0.0371 | 0.223 | 2.52 | 0.0139 | 0.16 |
| Q19749 | 0.242 | 0.508 | 0.734 | -0.201 | 0.58 | 0.789 |
| Q19766 | 0.0748 | 0.701 | 0.853 | 0.096 | 0.623 | 0.813 |
| Q19775 | 2.16 | 0.0073 | 0.121 | 0.0805 | 0.896 | 0.949 |
| Q19782 | 0.0294 | 0.943 | 0.971 | 0.969 | 0.0416 | 0.244 |
| Q19825 | 0.582 | 0.642 | 0.821 | -1.41 | 0.276 | 0.55 |
| Q19842 | 0.769 | 0.00892 | 0.128 | 0.443 | 0.0815 | 0.312 |
| Q19869 | -0.139 | 0.469 | 0.701 | -0.226 | 0.252 | 0.515 |
| Q19877 | 0.135 | 0.466 | 0.699 | -0.22 | 0.249 | 0.512 |
| Q19967 | 1.08 | 0.311 | 0.609 | 2.44 | 0.0417 | 0.244 |
| Q19969 | -0.694 | 0.466 | 0.699 | -3.11 | 0.00964 | 0.139 |
| Q19978 | 0.0998 | 0.941 | 0.971 | -3.84 | 0.0204 | 0.192 |
| Q20053 | 1.36 | 0.0867 | 0.331 | 1.09 | 0.154 | 0.407 |
| Q20140 | 0.459 | 0.66 | 0.831 | 1.63 | 0.144 | 0.399 |
| Q20168 | 0.44 | 0.636 | 0.819 | -0.822 | 0.384 | 0.65 |
| Q20222 | -0.0362 | 0.979 | 0.989 | -0.00433 | 0.998 | 0.998 |
| Q20224 | 0.579 | 0.313 | 0.612 | -0.129 | 0.816 | 0.913 |
| Q20228 | -0.342 | 0.215 | 0.514 | -0.402 | 0.153 | 0.407 |
| Q20334 | 0.323 | 0.385 | 0.659 | 0.496 | 0.196 | 0.456 |
| Q20363 | 0.637 | 0.0861 | 0.331 | 0.808 | 0.0383 | 0.237 |
| Q20448 | 0.16 | 0.64 | 0.821 | 0.349 | 0.321 | 0.598 |
| Q20496 | 1.63 | 0.00759 | 0.121 | 1.78 | 0.00485 | 0.1 |
| Q20507 | -1.43 | 0.237 | 0.538 | -2.13 | 0.0941 | 0.332 |
| Q20588 | -0.444 | 0.228 | 0.527 | -0.115 | 0.743 | 0.879 |
| Q20647 | 0.0356 | 0.861 | 0.936 | 0.0774 | 0.705 | 0.855 |
| Q20655 | 0.405 | 0.0652 | 0.293 | 0.106 | 0.588 | 0.791 |
| Q20719 | 1.28 | 0.158 | 0.434 | 0.862 | 0.323 | 0.6 |
| Q20728 | 0.0531 | 0.957 | 0.978 | 2.42 | 0.0355 | 0.232 |
| Q20748 | 0.884 | 0.349 | 0.637 | -1.43 | 0.148 | 0.4 |
| Q20751 | -0.307 | 0.199 | 0.492 | -0.0688 | 0.761 | 0.886 |
| Q20772 | 0.697 | 0.456 | 0.696 | 0.628 | 0.5 | 0.736 |
| Q20779 | -0.533 | 0.0876 | 0.332 | -0.279 | 0.336 | 0.612 |
| Q20898 | 0.378 | 0.8 | 0.909 | 0.678 | 0.651 | 0.831 |
| Q20938 | 0.882 | 0.562 | 0.774 | -0.0723 | 0.962 | 0.986 |
| Q20970 | 0.0661 | 0.94 | 0.971 | -1.23 | 0.189 | 0.448 |
| Q21067 | 0.926 | 0.323 | 0.617 | 2.52 | 0.0217 | 0.199 |
| Q21154 | -1.18 | 0.352 | 0.637 | 0.99 | 0.43 | 0.686 |
| Q21215 | -0.303 | 0.163 | 0.441 | -0.094 | 0.645 | 0.828 |
| Q21217 | 0.28 | 0.233 | 0.534 | -0.0876 | 0.696 | 0.854 |
| Q21265 | 0.365 | 0.181 | 0.464 | 0.772 | 0.0154 | 0.17 |
| Q21276 | 0.859 | 0.158 | 0.434 | 0.235 | 0.68 | 0.845 |
| Q21313 | 1.19 | 0.0292 | 0.212 | 0.592 | 0.221 | 0.481 |
| Q21351 | 0.233 | 0.304 | 0.604 | 0.241 | 0.287 | 0.565 |
| Q21443 | 0.25 | 0.322 | 0.616 | 0.512 | 0.0637 | 0.294 |
| Q21502 | 1.5 | 0.0646 | 0.292 | 1.05 | 0.171 | 0.425 |
| Q21551 | 0.195 | 0.439 | 0.685 | 0.597 | 0.038 | 0.236 |
| Q21568 | 0.999 | 0.107 | 0.361 | 0.799 | 0.184 | 0.443 |
| Q21633 | 0.21 | 0.678 | 0.843 | 0.673 | 0.207 | 0.47 |
| Q21693 | -0.281 | 0.642 | 0.821 | -0.583 | 0.347 | 0.622 |
| Q21735 | -0.125 | 0.89 | 0.941 | 0.189 | 0.835 | 0.92 |
| Q21752 | 0.236 | 0.269 | 0.571 | 0.266 | 0.217 | 0.479 |
| Q21824 | -0.225 | 0.309 | 0.608 | -0.163 | 0.454 | 0.703 |
| Q21832 | -0.132 | 0.811 | 0.911 | -0.275 | 0.621 | 0.813 |
| Q21926 | 1.38 | 0.354 | 0.638 | 0.133 | 0.926 | 0.965 |
| Q21930 | 0.0972 | 0.619 | 0.811 | 0.125 | 0.524 | 0.746 |
| Q21966 | 1.33 | 0.214 | 0.514 | 1.12 | 0.288 | 0.565 |
| Q21993 | 0.786 | 0.339 | 0.632 | 1.06 | 0.208 | 0.47 |
| Q22018 | -0.542 | 0.432 | 0.683 | -0.386 | 0.572 | 0.783 |
| Q22021 | 0.626 | 0.352 | 0.637 | 1.63 | 0.0337 | 0.232 |
| Q22037 | -0.149 | 0.644 | 0.821 | 0.141 | 0.661 | 0.837 |
| Q22038 | 0.529 | 0.0258 | 0.201 | 0.184 | 0.366 | 0.637 |
| Q22053 | 0.433 | 0.0988 | 0.349 | 0.18 | 0.457 | 0.703 |
| Q22054 | 0.0123 | 0.946 | 0.972 | -0.0793 | 0.664 | 0.838 |
| Q22067 | 1.12 | 0.0785 | 0.314 | 0.361 | 0.533 | 0.751 |
| Q22099 | 0.362 | 0.3 | 0.601 | -0.489 | 0.173 | 0.428 |
| Q22100 | 0.558 | 0.0777 | 0.314 | 0.586 | 0.0664 | 0.299 |
| Q22169 | -1.42 | 0.0726 | 0.302 | -1.73 | 0.0364 | 0.233 |
| Q22235 | 0.273 | 0.384 | 0.659 | 0.28 | 0.373 | 0.642 |
| Q22285 | 0.642 | 0.024 | 0.2 | 0.699 | 0.0165 | 0.179 |
| Q22288 | 0.165 | 0.582 | 0.787 | 0.329 | 0.285 | 0.562 |
| Q22347 | 0.0777 | 0.854 | 0.934 | 0.0015 | 0.997 | 0.998 |
| Q22494 | 0.765 | 0.535 | 0.756 | -0.633 | 0.606 | 0.802 |
| Q22505 | 0.333 | 0.293 | 0.595 | -0.0934 | 0.76 | 0.886 |
| Q22633 | -0.849 | 0.00244 | 0.0825 | -0.902 | 0.00172 | 0.0731 |
| Q22799 | 0.0636 | 0.876 | 0.94 | -0.693 | 0.119 | 0.366 |
| Q22866 | -0.865 | 0.0439 | 0.232 | -1.28 | 0.00798 | 0.132 |
| Q22957 | -0.188 | 0.836 | 0.923 | -0.169 | 0.853 | 0.928 |
| Q22993 | 0.41 | 0.181 | 0.464 | 0.461 | 0.138 | 0.387 |
| Q23120 | 0.0444 | 0.896 | 0.945 | 0.164 | 0.631 | 0.819 |
| Q23121 | -0.0148 | 0.941 | 0.971 | 0.139 | 0.496 | 0.731 |
| Q23280 | -0.598 | 0.498 | 0.724 | -0.866 | 0.334 | 0.61 |
| Q23307 | 0.281 | 0.339 | 0.632 | 0.49 | 0.115 | 0.36 |
| Q23312 | -0.248 | 0.229 | 0.529 | -0.49 | 0.0338 | 0.232 |
| Q23381 | 1.06 | 0.0956 | 0.345 | 1.15 | 0.0756 | 0.311 |
| Q23445 | -0.116 | 0.932 | 0.966 | 0.137 | 0.92 | 0.964 |
| Q23449 | -0.0226 | 0.979 | 0.989 | -0.187 | 0.828 | 0.917 |
| Q23500 | 0.136 | 0.489 | 0.719 | 0.167 | 0.4 | 0.66 |
| Q23551 | 0.348 | 0.249 | 0.547 | 0.227 | 0.441 | 0.692 |
| Q23588 | -0.388 | 0.287 | 0.587 | -0.121 | 0.73 | 0.873 |
| Q23655 | -0.42 | 0.24 | 0.538 | -0.59 | 0.112 | 0.358 |
| Q23670 | 1.3 | 0.345 | 0.636 | -1.15 | 0.401 | 0.66 |
| Q23680 | -0.418 | 0.243 | 0.543 | 0.247 | 0.477 | 0.712 |
| Q27245 | -1.61 | 0.00238 | 0.0825 | -0.58 | 0.148 | 0.4 |
| Q27249 | -0.597 | 0.134 | 0.401 | -0.967 | 0.0278 | 0.217 |
| Q27371 | -0.234 | 0.286 | 0.587 | -0.107 | 0.615 | 0.807 |
| Q27389 | -0.0616 | 0.742 | 0.875 | -0.195 | 0.313 | 0.591 |
| Q27487 | 2.19 | 0.188 | 0.474 | 1.41 | 0.382 | 0.65 |
| Q27488 | -0.862 | 0.285 | 0.587 | 0.331 | 0.671 | 0.839 |
| Q27504 | 0.513 | 0.382 | 0.659 | -2.13 | 0.00543 | 0.108 |
| Q27511 | 1.3 | 0.0256 | 0.201 | 1.09 | 0.0496 | 0.265 |
| Q27527 | -0.155 | 0.378 | 0.658 | 0.0831 | 0.63 | 0.819 |
| Q27535 | 0.713 | 0.0102 | 0.137 | 0.419 | 0.0832 | 0.316 |
| Q27876 | 0.992 | 0.693 | 0.849 | 1.67 | 0.51 | 0.741 |
| Q27888 | 0.195 | 0.807 | 0.911 | -0.0395 | 0.96 | 0.986 |
| Q27894 | 0.281 | 0.309 | 0.608 | -0.401 | 0.16 | 0.41 |
| Q4TT88 | 0.304 | 0.674 | 0.841 | -0.908 | 0.231 | 0.494 |
| Q564Q1 | 1.53 | 0.0495 | 0.251 | -0.039 | 0.954 | 0.984 |
| Q7JKP6 | 1.5 | 0.000802 | 0.0514 | -0.399 | 0.193 | 0.452 |
| Q7K6X4 | 1.33 | 0.0656 | 0.293 | 1.27 | 0.0767 | 0.311 |
| Q86DC6 | 0.774 | 0.0094 | 0.13 | 0.857 | 0.00561 | 0.11 |
| Q86S66 | -0.0523 | 0.811 | 0.911 | 0.0702 | 0.748 | 0.881 |
| Q8TA83 | -0.776 | 0.245 | 0.544 | -0.0392 | 0.951 | 0.983 |
| Q93235 | 0.443 | 0.159 | 0.434 | -0.185 | 0.534 | 0.751 |
| Q93244 | 0.241 | 0.598 | 0.798 | 0.904 | 0.0745 | 0.311 |
| Q93379 | -1.92 | 0.0718 | 0.3 | -2.29 | 0.0389 | 0.237 |
| Q93408 | -2.63 | 0.0602 | 0.283 | -0.874 | 0.485 | 0.72 |
| Q93561 | 1.06 | 0.223 | 0.521 | 1.55 | 0.0893 | 0.327 |
| Q93572 | -0.304 | 0.149 | 0.424 | -0.234 | 0.253 | 0.515 |
| Q93573 | -0.651 | 0.0295 | 0.212 | -0.285 | 0.276 | 0.55 |
| Q93615 | 0.0755 | 0.7 | 0.853 | 0.173 | 0.388 | 0.653 |
| Q93714 | 1.01 | 0.0013 | 0.0656 | 0.196 | 0.369 | 0.64 |
| Q93725 | 0.734 | 0.496 | 0.724 | 0.588 | 0.583 | 0.791 |
| Q93761 | 0.416 | 0.217 | 0.514 | 0.159 | 0.623 | 0.813 |
| Q94045 | 0.667 | 0.588 | 0.79 | 0.00668 | 0.996 | 0.998 |
| Q94051 | 0.394 | 0.822 | 0.916 | -0.0942 | 0.957 | 0.985 |
| Q94230 | 0.15 | 0.432 | 0.683 | 0.0475 | 0.801 | 0.902 |
| Q94234 | -1.19 | 0.245 | 0.544 | -1.5 | 0.151 | 0.404 |
| Q94261 | 0.379 | 0.752 | 0.88 | -1.95 | 0.133 | 0.382 |
| Q94272 | 0.375 | 0.0921 | 0.34 | 0.528 | 0.028 | 0.217 |
| Q94300 | 1.65 | 0.04 | 0.229 | 2.29 | 0.00969 | 0.139 |
| Q94360 | 1.36 | 0.00397 | 0.101 | -0.0401 | 0.908 | 0.955 |
| Q95005 | -0.25 | 0.361 | 0.645 | -0.417 | 0.145 | 0.4 |
| Q95008 | 1.28 | 0.434 | 0.683 | 0.521 | 0.745 | 0.879 |
| Q95017 | 2.39 | 0.0146 | 0.16 | 1.99 | 0.0316 | 0.228 |
| Q95QA6 | 0.387 | 0.155 | 0.434 | 0.525 | 0.0668 | 0.299 |
| Q95QV8 | 0.0695 | 0.926 | 0.964 | 0.459 | 0.543 | 0.76 |
| Q95QW0 | 0.58 | 0.282 | 0.584 | 0.175 | 0.736 | 0.877 |
| Q95X44 | 0.198 | 0.261 | 0.561 | 0.203 | 0.25 | 0.513 |
| Q95Y04 | -0.512 | 0.0466 | 0.24 | -0.0891 | 0.691 | 0.851 |
| Q95Y72 | 0.74 | 0.0411 | 0.229 | 1.39 | 0.00195 | 0.0743 |
| Q95Y90 | -0.257 | 0.317 | 0.612 | -0.189 | 0.455 | 0.703 |
| Q95YF3 | 0.196 | 0.604 | 0.803 | -0.243 | 0.521 | 0.744 |
| Q965S8 | 0.728 | 0.17 | 0.45 | 0.214 | 0.668 | 0.839 |
| Q966C6 | 0.165 | 0.35 | 0.637 | -0.188 | 0.293 | 0.571 |
| Q9BKS0 | 1.85 | 0.215 | 0.514 | 2.14 | 0.159 | 0.41 |
| Q9BKU4 | 0.567 | 0.286 | 0.587 | -0.732 | 0.179 | 0.436 |
| Q9BKU8 | -1.83 | 0.244 | 0.544 | -1.2 | 0.433 | 0.686 |
| Q9BL07 | 0.323 | 0.566 | 0.775 | -0.729 | 0.214 | 0.477 |
| Q9BL19 | -0.0321 | 0.849 | 0.93 | 0.051 | 0.763 | 0.886 |
| Q9GYF1 | 0.292 | 0.175 | 0.458 | 0.453 | 0.0508 | 0.268 |
| Q9GZE9 | -2.04 | 0.0122 | 0.151 | -0.8 | 0.237 | 0.501 |
| Q9GZH4 | 0.208 | 0.46 | 0.697 | -0.189 | 0.502 | 0.736 |
| Q9N2W5 | -0.0713 | 0.897 | 0.945 | 0.591 | 0.303 | 0.584 |
| Q9N2W7 | -0.157 | 0.867 | 0.939 | 0.562 | 0.554 | 0.771 |
| Q9N358 | 0.126 | 0.801 | 0.909 | -0.642 | 0.221 | 0.481 |
| Q9N3B0 | -1.52 | 0.374 | 0.654 | -1.17 | 0.49 | 0.726 |
| Q9N3X2 | -0.0672 | 0.728 | 0.869 | -0.532 | 0.0226 | 0.202 |
| Q9N408 | -0.447 | 0.182 | 0.466 | 0.421 | 0.206 | 0.47 |
| Q9N4A7 | 0.252 | 0.8 | 0.909 | -0.0505 | 0.959 | 0.986 |
| Q9N4D8 | -1.78 | 0.0506 | 0.254 | -1.28 | 0.136 | 0.385 |
| Q9N4G9 | 0.367 | 0.395 | 0.661 | 0.443 | 0.309 | 0.588 |
| Q9N4I4 | 0.155 | 0.454 | 0.696 | -0.081 | 0.691 | 0.851 |
| Q9N4J8 | 0.117 | 0.615 | 0.809 | -0.228 | 0.338 | 0.612 |
| Q9N4K0 | 2.72 | 0.121 | 0.381 | 2.39 | 0.165 | 0.416 |
| Q9N4M4 | 0.131 | 0.523 | 0.743 | 0.28 | 0.191 | 0.449 |
| Q9N4V0 | 1.93 | 0.13 | 0.395 | 4.12 | 0.00726 | 0.128 |
| Q9N599 | 0.199 | 0.693 | 0.849 | 0.266 | 0.599 | 0.798 |
| Q9N5M2 | -0.164 | 0.831 | 0.921 | -0.246 | 0.749 | 0.882 |
| Q9NAE2 | -0.606 | 0.524 | 0.744 | -1.5 | 0.14 | 0.389 |
| Q9NAP9 | 0.782 | 0.218 | 0.514 | 0.317 | 0.602 | 0.801 |
| Q9NEN6 | -0.117 | 0.641 | 0.821 | 0.173 | 0.493 | 0.729 |
| Q9NEU5 | -1.28 | 0.247 | 0.545 | -0.929 | 0.39 | 0.653 |
| Q9NEW6 | -0.151 | 0.476 | 0.708 | 0.00197 | 0.992 | 0.997 |
| Q9NEZ5 | -1.3 | 0.0179 | 0.18 | -1.36 | 0.0144 | 0.162 |
| Q9TXH9 | -0.931 | 0.319 | 0.615 | -2.85 | 0.0124 | 0.153 |
| Q9TXP0 | 0.0639 | 0.719 | 0.867 | 0.056 | 0.752 | 0.882 |
| Q9TYK1 | 1.5 | 0.0361 | 0.221 | 1.93 | 0.0122 | 0.153 |
| Q9TYV5 | 1.4 | 0.0304 | 0.212 | 1.12 | 0.0687 | 0.301 |
| Q9TYW1 | 0.779 | 0.424 | 0.679 | -0.622 | 0.52 | 0.744 |
| Q9TZQ3 | -0.0602 | 0.95 | 0.973 | -0.464 | 0.634 | 0.821 |
| Q9U1Q4 | 1.75 | 0.107 | 0.361 | 0.579 | 0.564 | 0.779 |
| Q9U1W1 | -0.289 | 0.387 | 0.66 | 0.255 | 0.443 | 0.693 |
| Q9U256 | -1.44 | 0.0999 | 0.349 | 0.578 | 0.474 | 0.712 |
| Q9U2A8 | -0.019 | 0.934 | 0.966 | 0.133 | 0.566 | 0.78 |
| Q9U2D9 | -0.202 | 0.802 | 0.909 | -0.862 | 0.302 | 0.584 |
| Q9U2H9 | -0.0882 | 0.69 | 0.849 | 0.149 | 0.506 | 0.738 |
| Q9U2X0 | -0.212 | 0.685 | 0.845 | -1.27 | 0.0371 | 0.235 |
| Q9U332 | -0.0681 | 0.772 | 0.894 | -0.17 | 0.476 | 0.712 |
| Q9U3F4 | -0.038 | 0.894 | 0.943 | 0.225 | 0.439 | 0.691 |
| Q9UAV5 | 0.611 | 0.0126 | 0.151 | 0.244 | 0.233 | 0.497 |
| Q9XTI0 | 0.516 | 0.619 | 0.811 | 1.27 | 0.241 | 0.504 |
| Q9XTT9 | 1.55 | 0.375 | 0.654 | 2.32 | 0.197 | 0.457 |
| Q9XUN9 | -0.565 | 0.728 | 0.869 | -0.335 | 0.836 | 0.921 |
| Q9XUV0 | 1.25 | 0.0603 | 0.283 | 1.27 | 0.0561 | 0.277 |
| Q9XUY5 | 1.68 | 0.0997 | 0.349 | 1.72 | 0.0931 | 0.331 |
| Q9XVF7 | 0.176 | 0.326 | 0.617 | 0.209 | 0.25 | 0.513 |
| Q9XVI9 | -0.554 | 0.448 | 0.694 | -1.01 | 0.185 | 0.444 |
| Q9XVP0 | -0.0236 | 0.914 | 0.956 | -0.425 | 0.0803 | 0.312 |
| Q9XVR8 | 0.178 | 0.644 | 0.821 | 0.421 | 0.289 | 0.565 |
| Q9XVS4 | 0.8 | 0.0463 | 0.24 | 1.34 | 0.00452 | 0.0997 |
| Q9XVT0 | 1.37 | 0.191 | 0.48 | 1.85 | 0.0901 | 0.328 |
| Q9XW16 | -0.0292 | 0.887 | 0.941 | -0.245 | 0.256 | 0.52 |
| Q9XW17 | 0.326 | 0.515 | 0.737 | 0.13 | 0.793 | 0.9 |
| Q9XW92 | 0.302 | 0.245 | 0.544 | -0.0632 | 0.799 | 0.902 |
| Q9XWI6 | 0.456 | 0.368 | 0.651 | -0.592 | 0.251 | 0.514 |
| Q9XWN7 | -1.38 | 0.0394 | 0.228 | -0.174 | 0.763 | 0.886 |
| Q9XWU2 | 1.45 | 0.128 | 0.395 | 1.87 | 0.0606 | 0.292 |
| Q9XWV2 | -0.75 | 0.352 | 0.637 | -1.66 | 0.061 | 0.292 |
| Q9XXD4 | -1.37 | 0.384 | 0.659 | 0.335 | 0.828 | 0.917 |
| Q9XXK1 | 0.109 | 0.555 | 0.771 | -0.244 | 0.208 | 0.47 |
| Q9XXU9 | 0.253 | 0.366 | 0.648 | 0.242 | 0.385 | 0.65 |
| Q9Y0V6 | -0.144 | 0.666 | 0.836 | 0.839 | 0.0322 | 0.228 |
| V6CLP5 | 0.644 | 0.0317 | 0.213 | 0.268 | 0.308 | 0.588 |
| A0A061ACP8 | 0.274 | 0.315 | 0.612 | 0.342 | 0.219 | 0.48 |
| A0A061AE99 | -0.098 | 0.839 | 0.924 | 0.574 | 0.255 | 0.52 |
| A0A061AKY5 | 1.49 | 0.036 | 0.221 | 1.74 | 0.0191 | 0.189 |
| A0A078BS22 | 1.29 | 0.144 | 0.417 | 2.44 | 0.0161 | 0.176 |
| A0A0K3AQS9 | -0.772 | 0.0344 | 0.217 | -0.851 | 0.023 | 0.202 |
| A0A0K3ASN4 | 0.687 | 0.0612 | 0.285 | -0.0464 | 0.886 | 0.945 |
| A0A0K3AT05 | -1.35 | 0.0313 | 0.212 | -0.00617 | 0.991 | 0.997 |
| A0A0K3AVF2 | 0.278 | 0.781 | 0.902 | 2.03 | 0.0699 | 0.302 |
| A0A0K3AVS5 | -0.118 | 0.87 | 0.939 | 0.73 | 0.325 | 0.602 |
| A0A0K3AXM4 | -2.69 | 0.0251 | 0.2 | -1.23 | 0.24 | 0.504 |
| A0A0K3AYJ1 | 0.5 | 0.0667 | 0.294 | 0.699 | 0.0184 | 0.188 |
| A0A0S4XR43 | -3.75 | 0.173 | 0.457 | -2.09 | 0.426 | 0.684 |
| A0A164D3G3 | 1.25 | 0.0165 | 0.171 | 1.93 | 0.00172 | 0.0731 |
| A0A1I6CMC9 | -1.12 | 0.124 | 0.387 | 0.0973 | 0.884 | 0.944 |
| A0A1N7SYN7 | 0.456 | 0.747 | 0.878 | 0.36 | 0.798 | 0.902 |
| A0A2C9C377 | 0.933 | 0.42 | 0.676 | 1.89 | 0.124 | 0.372 |
| A0A2C9C3A0 | -1.66 | 0.129 | 0.395 | -2.47 | 0.0362 | 0.233 |
| A0A2C9C3S7 | 1.71 | 0.00797 | 0.121 | 0.613 | 0.239 | 0.503 |
| A0A2K5ATV3 | -0.569 | 0.488 | 0.719 | 0.172 | 0.832 | 0.919 |
| A0A486WV52 | 0.625 | 0.0739 | 0.305 | 0.577 | 0.0941 | 0.332 |
| A0A486WWU7 | 0.802 | 0.0339 | 0.217 | 1.06 | 0.01 | 0.139 |
| A0A486WX07 | 0.165 | 0.811 | 0.911 | -2.48 | 0.00634 | 0.117 |
| A0A486WX27 | 0.746 | 0.375 | 0.654 | 0.303 | 0.712 | 0.859 |
| A0A486WXP6 | 0.543 | 0.193 | 0.481 | 0.481 | 0.243 | 0.504 |
| A0A4V0IIR6 | -0.135 | 0.451 | 0.696 | -0.116 | 0.515 | 0.742 |
| A0A4V0IJD5 | -0.0851 | 0.807 | 0.911 | -0.513 | 0.168 | 0.42 |
| A0A5E4LWV7 | 0.336 | 0.183 | 0.467 | 0.195 | 0.421 | 0.679 |
| A0A5S9MMA9 | -0.458 | 0.732 | 0.869 | -1.39 | 0.314 | 0.592 |
| A0A7R9XLP7 | -1.47 | 0.0172 | 0.175 | -2.1 | 0.00284 | 0.0822 |
| A0A8S4QD99 | 1.43 | 0.162 | 0.438 | -1.26 | 0.21 | 0.471 |
| A0A9J5DXX3 | -2.22 | 0.0512 | 0.256 | -2.28 | 0.0469 | 0.258 |
| A0A9J5HVW4 | 0.781 | 0.175 | 0.458 | 0.531 | 0.341 | 0.616 |
| A3QM98 | -3.97 | 0.0166 | 0.171 | -3.59 | 0.0255 | 0.216 |
| A3QMC5 | -0.162 | 0.392 | 0.661 | -0.0726 | 0.696 | 0.854 |
| A5Z2T8 | 1.57 | 0.184 | 0.469 | 2.21 | 0.0759 | 0.311 |
| A6ZJ46 | 1.39 | 0.263 | 0.564 | 0.217 | 0.855 | 0.93 |
| A7DT45 | -0.86 | 0.066 | 0.293 | -1.29 | 0.0129 | 0.153 |
| A7DTF5 | 0.395 | 0.454 | 0.696 | 0.221 | 0.671 | 0.839 |
| A8WI97 | 0.104 | 0.917 | 0.958 | 0.525 | 0.604 | 0.802 |
| A9UJN7 | 0.191 | 0.839 | 0.924 | -1.14 | 0.249 | 0.512 |
| B0M0L8 | 0.0572 | 0.917 | 0.958 | 1.13 | 0.0677 | 0.301 |
| B1Q273 | -0.463 | 0.148 | 0.423 | 0.514 | 0.114 | 0.36 |
| B2D6P1 | 0.107 | 0.706 | 0.855 | 0.217 | 0.452 | 0.702 |
| B3CJ34 | 0.582 | 0.503 | 0.729 | 1.3 | 0.157 | 0.41 |
| B3GWA1 | 1.54 | 0.0197 | 0.186 | 2.46 | 0.0018 | 0.0731 |
| B5U8N2 | 0.424 | 0.204 | 0.499 | 0.294 | 0.365 | 0.635 |
| B6VQ96 | -0.102 | 0.888 | 0.941 | 0.249 | 0.732 | 0.874 |
| B7CED8 | 0.939 | 0.0169 | 0.173 | -0.485 | 0.156 | 0.409 |
| B7FAR9 | 0.164 | 0.662 | 0.832 | 0.854 | 0.0472 | 0.259 |
| B7WN91 | 2.15 | 0.0382 | 0.226 | 1.59 | 0.103 | 0.341 |
| B9WRT9 | 0.53 | 0.087 | 0.331 | 0.608 | 0.0559 | 0.277 |
| C1P636 | 0.727 | 0.064 | 0.292 | 0.655 | 0.0888 | 0.327 |
| C6KRI9 | -2.43 | 0.108 | 0.361 | -0.194 | 0.888 | 0.946 |
| C6KRN4 | -1.23 | 0.101 | 0.349 | -1.24 | 0.0988 | 0.337 |
| C9IY22 | 1.06 | 0.0677 | 0.294 | 0.61 | 0.256 | 0.52 |
| D0IMZ5 | 0.524 | 0.0338 | 0.217 | 0.285 | 0.199 | 0.458 |
| D5MCQ2 | 0.583 | 0.0705 | 0.299 | -0.0959 | 0.739 | 0.878 |
| D5MCR9 | -0.452 | 0.742 | 0.875 | -1.84 | 0.205 | 0.469 |
| D7SFI3 | -1.07 | 0.00269 | 0.0841 | -0.13 | 0.61 | 0.806 |
| E3W741 | 0.634 | 0.0471 | 0.242 | 0.642 | 0.0449 | 0.254 |
| G4S034 | 0.596 | 0.0386 | 0.226 | 0.698 | 0.0203 | 0.192 |
| G4SI07 | -0.0062 | 0.995 | 0.998 | -1.05 | 0.319 | 0.598 |
| G4SJZ3 | -1.51 | 0.0329 | 0.214 | -0.79 | 0.213 | 0.476 |
| G4SL51 | 2.36 | 0.00558 | 0.109 | 1.61 | 0.0321 | 0.228 |
| G5EBF3 | -0.851 | 0.00602 | 0.109 | -0.623 | 0.026 | 0.216 |
| G5EBH7 | -1.42 | 0.013 | 0.151 | -0.981 | 0.0581 | 0.287 |
| G5EBI0 | -3.16 | 0.00395 | 0.101 | -2.64 | 0.01 | 0.139 |
| G5EBJ7 | -0.694 | 0.0495 | 0.251 | -1.18 | 0.00454 | 0.0997 |
| G5EBK3 | 0.198 | 0.683 | 0.845 | -0.0302 | 0.95 | 0.983 |
| G5EBR1 | 0.0939 | 0.732 | 0.869 | -0.0606 | 0.824 | 0.917 |
| G5EBY6 | -2.29 | 0.0187 | 0.184 | -2.75 | 0.00783 | 0.132 |
| G5EC10 | -1.47 | 0.369 | 0.652 | -1.42 | 0.385 | 0.65 |
| G5EC31 | -0.111 | 0.849 | 0.93 | -0.961 | 0.128 | 0.375 |
| G5EC65 | 2.19 | 0.00272 | 0.0841 | 1.91 | 0.0058 | 0.112 |
| G5EC71 | -0.123 | 0.614 | 0.809 | 0.162 | 0.509 | 0.741 |
| G5EC87 | -0.136 | 0.741 | 0.875 | -0.766 | 0.0918 | 0.33 |
| G5ECA7 | 0.324 | 0.634 | 0.819 | 0.602 | 0.384 | 0.65 |
| G5ECE7 | -1.03 | 0.298 | 0.601 | -0.225 | 0.814 | 0.911 |
| G5ECG8 | 0.399 | 0.087 | 0.331 | 0.11 | 0.604 | 0.802 |
| G5ECL3 | 0.378 | 0.622 | 0.812 | -1.45 | 0.087 | 0.324 |
| G5ECM6 | 0.0737 | 0.731 | 0.869 | -0.031 | 0.884 | 0.944 |
| G5ECR0 | 0.693 | 0.186 | 0.471 | 0.0196 | 0.968 | 0.986 |
| G5ECR7 | -1.1 | 0.287 | 0.587 | -1.57 | 0.143 | 0.395 |
| G5ECS9 | 0.826 | 0.374 | 0.654 | -0.926 | 0.322 | 0.598 |
| G5ECT6 | -0.136 | 0.717 | 0.867 | 0.0364 | 0.922 | 0.964 |
| G5ECV9 | 2.45 | 0.0298 | 0.212 | 2.01 | 0.0615 | 0.294 |
| G5ED01 | -0.349 | 0.51 | 0.734 | -2.11 | 0.00348 | 0.0942 |
| G5ED07 | -0.263 | 0.199 | 0.492 | 0.0287 | 0.882 | 0.943 |
| G5ED31 | 0.0507 | 0.826 | 0.917 | -0.0933 | 0.687 | 0.848 |
| G5EDD4 | 0.172 | 0.685 | 0.845 | -0.397 | 0.362 | 0.632 |
| G5EDT4 | -1.18 | 0.0828 | 0.324 | 0.403 | 0.516 | 0.742 |
| G5EDV3 | 0.427 | 0.185 | 0.47 | 0.948 | 0.0128 | 0.153 |
| G5EDW8 | -0.768 | 0.129 | 0.395 | 0.657 | 0.185 | 0.444 |
| G5EDZ5 | 1.31 | 0.12 | 0.379 | 2.05 | 0.0268 | 0.217 |
| G5EE04 | -0.272 | 0.471 | 0.703 | -0.334 | 0.379 | 0.647 |
| G5EE41 | -2.6 | 0.00132 | 0.0656 | -2.83 | 0.000796 | 0.0605 |
| G5EE74 | 1.53 | 0.235 | 0.538 | 1.61 | 0.214 | 0.477 |
| G5EE90 | -0.448 | 0.721 | 0.868 | -0.414 | 0.742 | 0.879 |
| G5EEA8 | 0.253 | 0.451 | 0.696 | -0.24 | 0.473 | 0.712 |
| G5EEG4 | 0.621 | 0.28 | 0.583 | 0.845 | 0.154 | 0.407 |
| G5EEG8 | 1.14 | 0.00434 | 0.106 | 0.913 | 0.0132 | 0.155 |
| G5EEK8 | 0.713 | 0.0325 | 0.214 | 0.652 | 0.0456 | 0.256 |
| G5EEL9 | -0.013 | 0.969 | 0.984 | -0.916 | 0.0248 | 0.214 |
| G5EET8 | -1.02 | 0.00107 | 0.0589 | -0.565 | 0.0228 | 0.202 |
| G5EEV5 | 0.197 | 0.807 | 0.911 | 0.0357 | 0.965 | 0.986 |
| G5EF32 | -0.788 | 0.0278 | 0.21 | -0.623 | 0.0662 | 0.299 |
| G5EF37 | -0.795 | 0.0815 | 0.32 | 0.189 | 0.647 | 0.83 |
| G5EF97 | -0.0617 | 0.908 | 0.951 | 0.21 | 0.695 | 0.854 |
| G5EFE3 | 0.446 | 0.119 | 0.378 | 0.749 | 0.0194 | 0.189 |
| G5EFG4 | 0.81 | 0.465 | 0.699 | 0.498 | 0.65 | 0.831 |
| G5EFI4 | 0.319 | 0.523 | 0.743 | -0.525 | 0.304 | 0.584 |
| G5EFL5 | 0.0119 | 0.965 | 0.982 | -0.274 | 0.331 | 0.608 |
| G5EFM6 | 0.684 | 0.135 | 0.401 | 0.866 | 0.0689 | 0.301 |
| G5EFP2 | 0.158 | 0.765 | 0.891 | 0.347 | 0.515 | 0.742 |
| G5EFS5 | 0.859 | 0.0283 | 0.212 | 0.0547 | 0.868 | 0.933 |
| G5EFV4 | 0.816 | 0.00356 | 0.0995 | 0.853 | 0.00278 | 0.0822 |
| G5EG13 | 1.36 | 0.117 | 0.376 | 2.19 | 0.023 | 0.202 |
| G5EGB1 | 0.813 | 0.0156 | 0.165 | 0.653 | 0.039 | 0.237 |
| G5EGK1 | -0.665 | 0.651 | 0.826 | -1.06 | 0.476 | 0.712 |
| H2FLK8 | -0.143 | 0.731 | 0.869 | -0.302 | 0.476 | 0.712 |
| H2KY95 | -1.88 | 0.418 | 0.674 | -3.65 | 0.137 | 0.385 |
| H2KYI5 | 0.066 | 0.879 | 0.94 | 0.236 | 0.588 | 0.791 |
| H2KYJ5 | -1.09 | 0.233 | 0.534 | -0.531 | 0.548 | 0.765 |
| H2KYK9 | -1.29 | 0.0422 | 0.23 | -1.77 | 0.0108 | 0.141 |
| H2KYP5 | -2.89 | 0.258 | 0.558 | -4.06 | 0.126 | 0.372 |
| H2KYR1 | 0.0724 | 0.733 | 0.869 | 0.0292 | 0.89 | 0.948 |
| H2KYV3 | 0.53 | 0.0754 | 0.31 | 0.119 | 0.657 | 0.835 |
| H2KZ73 | -0.178 | 0.87 | 0.939 | 1.75 | 0.136 | 0.385 |
| H2KZS2 | -0.0159 | 0.947 | 0.972 | 0.413 | 0.115 | 0.36 |
| H2KZV5 | -0.0186 | 0.928 | 0.965 | 0.312 | 0.158 | 0.41 |
| H2KZV8 | -0.12 | 0.743 | 0.876 | 0.672 | 0.0946 | 0.333 |
| H2KZY6 | 1.61 | 0.153 | 0.431 | 0.694 | 0.514 | 0.742 |
| H2L023 | 0.293 | 0.785 | 0.904 | 0.633 | 0.56 | 0.777 |
| H2L044 | -0.576 | 0.459 | 0.697 | -0.67 | 0.392 | 0.653 |
| H2L048 | 0.48 | 0.105 | 0.36 | -0.262 | 0.347 | 0.622 |
| H2L0F5 | 1.76 | 0.16 | 0.436 | -0.904 | 0.449 | 0.699 |
| H2L0G8 | 0.6 | 0.379 | 0.658 | 3.19 | 0.00128 | 0.0731 |
| H2L0Q5 | -1.58 | 0.274 | 0.576 | 2.75 | 0.0766 | 0.311 |
| H2L0S8 | -0.108 | 0.844 | 0.927 | 0.882 | 0.135 | 0.385 |
| H2L2A0 | -0.184 | 0.749 | 0.878 | -1.01 | 0.106 | 0.347 |
| H2L2C9 | 0.151 | 0.629 | 0.813 | 0.565 | 0.098 | 0.337 |
| H2L2E8 | 0.116 | 0.593 | 0.793 | -0.204 | 0.357 | 0.63 |
| H9G2T4 | 0.429 | 0.0366 | 0.221 | 0.346 | 0.0771 | 0.311 |
| I2HAJ8 | 0.604 | 0.228 | 0.527 | 0.202 | 0.673 | 0.839 |
| K8ERV5 | 0.152 | 0.685 | 0.845 | 0.232 | 0.54 | 0.757 |
| K8ESM2 | 0.939 | 0.443 | 0.69 | 1.21 | 0.331 | 0.608 |
| K8FDY0 | 0.0693 | 0.874 | 0.939 | 0.67 | 0.153 | 0.407 |
| L8E833 | -0.267 | 0.511 | 0.735 | -0.628 | 0.146 | 0.4 |
| M1ZJ62 | 0.262 | 0.399 | 0.662 | -0.0484 | 0.873 | 0.937 |
| O01497 | -0.603 | 0.374 | 0.654 | -0.793 | 0.251 | 0.514 |
| O01542 | 0.344 | 0.0937 | 0.342 | 0.246 | 0.209 | 0.47 |
| O01593 | -0.0619 | 0.947 | 0.972 | -0.305 | 0.745 | 0.879 |
| O01685 | 1.12 | 0.00672 | 0.117 | 0.451 | 0.176 | 0.432 |
| O01780 | 1.54 | 0.0303 | 0.212 | 1.1 | 0.0967 | 0.336 |
| O01806 | 0.893 | 0.00864 | 0.125 | 1.03 | 0.00416 | 0.0992 |
| O01816 | -0.483 | 0.46 | 0.697 | -0.902 | 0.186 | 0.445 |
| O01869 | 0.409 | 0.119 | 0.379 | 0.509 | 0.0619 | 0.294 |
| O01964 | -1.25 | 0.316 | 0.612 | 0.0456 | 0.97 | 0.986 |
| O02141 | 1.17 | 0.249 | 0.547 | 1.76 | 0.0997 | 0.337 |
| O02155 | 1.01 | 0.0209 | 0.189 | 0.781 | 0.056 | 0.277 |
| O02158 | 0.597 | 0.229 | 0.528 | 0.905 | 0.0842 | 0.318 |
| O02252 | 0.693 | 0.477 | 0.708 | -0.726 | 0.457 | 0.703 |
| O02267 | -0.28 | 0.401 | 0.662 | 0.0096 | 0.976 | 0.986 |
| O02286 | 0.352 | 0.157 | 0.434 | -0.0715 | 0.758 | 0.886 |
| O02357 | -3.72 | 0.00495 | 0.107 | -0.849 | 0.399 | 0.659 |
| O02637 | 0.58 | 0.268 | 0.571 | 1.21 | 0.0386 | 0.237 |
| O02642 | 0.519 | 0.13 | 0.395 | -0.0808 | 0.799 | 0.902 |
| O16226 | 0.993 | 0.101 | 0.349 | 1.24 | 0.0502 | 0.267 |
| O16249 | 1.21 | 0.029 | 0.212 | -0.412 | 0.389 | 0.653 |
| O16266 | 0.252 | 0.637 | 0.821 | -0.246 | 0.645 | 0.828 |
| O16298 | -0.157 | 0.567 | 0.775 | 0.129 | 0.639 | 0.826 |
| O16303 | -0.847 | 0.399 | 0.662 | 0.118 | 0.904 | 0.952 |
| O16309 | -2.85 | 0.112 | 0.368 | -2.88 | 0.109 | 0.351 |
| O16517 | -1.14 | 0.00682 | 0.117 | -1.12 | 0.00741 | 0.129 |
| O16521 | -0.0243 | 0.961 | 0.979 | -0.432 | 0.392 | 0.653 |
| O16997 | 1.55 | 0.217 | 0.514 | 2.18 | 0.0978 | 0.337 |
| O17218 | -0.167 | 0.576 | 0.781 | -0.115 | 0.7 | 0.854 |
| O17328 | -1.96 | 0.0405 | 0.229 | -1.7 | 0.0665 | 0.299 |
| O17345 | 0.667 | 0.209 | 0.508 | 0.427 | 0.406 | 0.664 |
| O17352 | -2.28 | 0.138 | 0.407 | -1.36 | 0.352 | 0.625 |
| O17406 | 0.284 | 0.293 | 0.595 | 0.633 | 0.0374 | 0.235 |
| O17626 | -0.289 | 0.802 | 0.909 | 0.0493 | 0.966 | 0.986 |
| O17643 | -0.153 | 0.882 | 0.941 | -0.4 | 0.7 | 0.854 |
| O17687 | 0.352 | 0.414 | 0.671 | 0.22 | 0.605 | 0.802 |
| O17694 | 0.272 | 0.36 | 0.645 | 0.0506 | 0.861 | 0.931 |
| O17725 | 0.982 | 0.344 | 0.636 | -2.11 | 0.0635 | 0.294 |
| O17759 | 0.297 | 0.236 | 0.538 | 0.00548 | 0.982 | 0.99 |
| O17836 | -0.142 | 0.615 | 0.809 | 0.117 | 0.676 | 0.841 |
| O17891 | -0.696 | 0.544 | 0.762 | -1.58 | 0.19 | 0.448 |
| O17892 | -0.255 | 0.76 | 0.888 | 0.109 | 0.896 | 0.949 |
| O17921 | -0.146 | 0.658 | 0.831 | -0.209 | 0.53 | 0.748 |
| O18000 | -1.57 | 0.202 | 0.498 | -0.371 | 0.751 | 0.882 |
| O18089 | 0.869 | 0.316 | 0.612 | -0.225 | 0.788 | 0.898 |
| O18180 | -0.378 | 0.436 | 0.683 | -0.184 | 0.699 | 0.854 |
| O18181 | 0.894 | 0.043 | 0.231 | 0.914 | 0.0398 | 0.237 |
| O18236 | 1.31 | 0.118 | 0.378 | 1.3 | 0.121 | 0.367 |
| O18239 | -2.34 | 0.0234 | 0.199 | -1.49 | 0.112 | 0.358 |
| O18240 | -0.198 | 0.3 | 0.601 | -0.232 | 0.231 | 0.494 |
| O44144 | 0.153 | 0.61 | 0.808 | 0.109 | 0.714 | 0.859 |
| O44145 | -0.0673 | 0.872 | 0.939 | 0.232 | 0.581 | 0.79 |
| O44444 | 0.255 | 0.424 | 0.679 | 0.647 | 0.0663 | 0.299 |
| O44471 | -0.366 | 0.682 | 0.845 | 0.605 | 0.502 | 0.736 |
| O44503 | 0.88 | 0.324 | 0.617 | -0.567 | 0.517 | 0.743 |
| O44509 | 0.976 | 0.304 | 0.604 | 0.748 | 0.424 | 0.68 |
| O44512 | -0.0762 | 0.702 | 0.853 | -0.198 | 0.332 | 0.609 |
| O44555 | -0.867 | 0.382 | 0.659 | -0.559 | 0.567 | 0.781 |
| O44565 | 3.19 | 0.0306 | 0.212 | 3.03 | 0.0374 | 0.235 |
| O44727 | -0.551 | 0.496 | 0.724 | 1.36 | 0.117 | 0.362 |
| O44772 | 1.13 | 0.109 | 0.363 | 0.104 | 0.872 | 0.937 |
| O44895 | 0.588 | 0.66 | 0.831 | -0.557 | 0.677 | 0.841 |
| O44906 | 1.22 | 0.00593 | 0.109 | 0.698 | 0.0644 | 0.296 |
| O44995 | 1.44 | 0.255 | 0.554 | 0.496 | 0.683 | 0.847 |
| O45011 | -0.27 | 0.411 | 0.67 | -0.604 | 0.0892 | 0.327 |
| O45012 | 0.666 | 0.0118 | 0.148 | 0.406 | 0.0815 | 0.312 |
| O45060 | -1.1 | 0.00192 | 0.0802 | -0.81 | 0.00978 | 0.139 |
| O45097 | -0.718 | 0.587 | 0.79 | -0.121 | 0.926 | 0.965 |
| O45106 | 1.01 | 0.226 | 0.527 | 0.183 | 0.818 | 0.913 |
| O45148 | -0.135 | 0.799 | 0.909 | -0.274 | 0.609 | 0.805 |
| O45226 | 0.904 | 0.373 | 0.654 | -0.847 | 0.403 | 0.661 |
| O45418 | 0.335 | 0.24 | 0.538 | 0.885 | 0.0107 | 0.141 |
| O45430 | 1.32 | 0.273 | 0.576 | 2.43 | 0.0634 | 0.294 |
| O45444 | -0.116 | 0.62 | 0.811 | 0.351 | 0.16 | 0.41 |
| O45451 | -1.8 | 0.0942 | 0.343 | -0.929 | 0.353 | 0.625 |
| O45502 | 0.471 | 0.525 | 0.744 | -1.44 | 0.0778 | 0.311 |
| O45525 | 0.89 | 0.394 | 0.661 | 1.24 | 0.244 | 0.506 |
| O45552 | -0.0703 | 0.725 | 0.869 | -0.179 | 0.381 | 0.649 |
| O45622 | 1.17 | 0.044 | 0.232 | 0.619 | 0.24 | 0.504 |
| O45712 | -0.563 | 0.405 | 0.666 | 0.918 | 0.191 | 0.449 |
| O45713 | -0.0288 | 0.949 | 0.972 | 0.239 | 0.597 | 0.798 |
| O45781 | 0.793 | 0.0925 | 0.34 | 1.54 | 0.00635 | 0.117 |
| O45783 | 3.25 | 0.00763 | 0.121 | 4.07 | 0.00227 | 0.0758 |
| O45815 | -0.0983 | 0.679 | 0.843 | -0.529 | 0.0514 | 0.269 |
| O45864 | -0.0987 | 0.65 | 0.825 | 0.358 | 0.127 | 0.372 |
| O45865 | -0.123 | 0.629 | 0.813 | -0.282 | 0.284 | 0.561 |
| O45934 | 2.3 | 0.0291 | 0.212 | 1.61 | 0.0985 | 0.337 |
| O45948 | 3.23 | 0.0419 | 0.23 | 5.82 | 0.00258 | 0.0786 |
| O61217 | 0.429 | 0.514 | 0.737 | 1.71 | 0.0276 | 0.217 |
| O61792 | 0.955 | 0.461 | 0.698 | 2.17 | 0.117 | 0.362 |
| O61793 | -3.79 | 0.00792 | 0.121 | -2.57 | 0.0433 | 0.249 |
| O61848 | -0.178 | 0.726 | 0.869 | 0.404 | 0.433 | 0.686 |
| O61880 | 0.23 | 0.474 | 0.707 | 0.404 | 0.224 | 0.486 |
| O62102 | -0.117 | 0.873 | 0.939 | -0.94 | 0.224 | 0.486 |
| O62198 | 1.69 | 0.0201 | 0.186 | 1.44 | 0.0393 | 0.237 |
| O62213 | -0.31 | 0.291 | 0.594 | -0.256 | 0.377 | 0.645 |
| O62277 | -0.677 | 0.0434 | 0.231 | -0.57 | 0.078 | 0.311 |
| O62289 | -0.233 | 0.771 | 0.894 | -2.6 | 0.0107 | 0.141 |
| O62337 | -0.311 | 0.321 | 0.616 | -0.0658 | 0.829 | 0.917 |
| O62388 | -0.17 | 0.563 | 0.774 | 0.569 | 0.0798 | 0.312 |
| O76367 | 0.983 | 0.0236 | 0.199 | 1.17 | 0.0109 | 0.141 |
| O76672 | 0.0827 | 0.941 | 0.971 | 0.257 | 0.818 | 0.913 |
| O76836 | 1.57 | 0.058 | 0.278 | 1.26 | 0.115 | 0.36 |
| P90732 | -0.232 | 0.73 | 0.869 | -1.08 | 0.135 | 0.385 |
| P90735 | 0.526 | 0.0565 | 0.273 | 0.406 | 0.123 | 0.372 |
| P90779 | -2.74 | 0.0209 | 0.189 | -1.86 | 0.0849 | 0.32 |
| P90849 | 0.00304 | 0.992 | 0.998 | 0.394 | 0.244 | 0.506 |
| P90860 | 0.135 | 0.583 | 0.787 | -0.924 | 0.00486 | 0.1 |
| P90867 | 0.884 | 0.248 | 0.545 | 1.01 | 0.192 | 0.45 |
| P90868 | 0.981 | 0.216 | 0.514 | 1.13 | 0.16 | 0.41 |
| P90889 | 0.146 | 0.784 | 0.904 | 0.443 | 0.415 | 0.674 |
| P90961 | -0.907 | 0.00599 | 0.109 | -0.374 | 0.16 | 0.41 |
| P90983 | 0.585 | 0.0676 | 0.294 | 0.664 | 0.0435 | 0.249 |
| P91020 | -0.279 | 0.252 | 0.552 | 0.305 | 0.215 | 0.477 |
| P91156 | -0.505 | 0.612 | 0.809 | -0.43 | 0.665 | 0.838 |
| P91180 | -1.41 | 0.26 | 0.56 | -2.37 | 0.076 | 0.311 |
| P91207 | -3.42 | 0.00578 | 0.109 | -3.64 | 0.00418 | 0.0992 |
| P91250 | -2.5 | 0.0491 | 0.251 | -2.69 | 0.0376 | 0.235 |
| P91283 | 0.201 | 0.297 | 0.601 | -0.265 | 0.18 | 0.436 |
| P91306 | 0.282 | 0.659 | 0.831 | 0.166 | 0.795 | 0.901 |
| P91398 | 0.957 | 0.259 | 0.559 | 0.691 | 0.406 | 0.663 |
| P91423 | -0.474 | 0.0553 | 0.27 | 0.0524 | 0.809 | 0.907 |
| P91453 | 1.22 | 0.363 | 0.646 | 1.82 | 0.189 | 0.448 |
| P91457 | -0.266 | 0.802 | 0.909 | -0.0363 | 0.973 | 0.986 |
| P92005 | 0.998 | 0.0895 | 0.336 | 1.32 | 0.0346 | 0.232 |
| Q09489 | -0.358 | 0.339 | 0.632 | -0.0911 | 0.802 | 0.902 |
| Q09567 | 1.01 | 0.0607 | 0.284 | 1.61 | 0.00884 | 0.134 |
| Q09979 | 2.16 | 0.0108 | 0.138 | 1.66 | 0.0341 | 0.232 |
| Q0G820 | 1.04 | 0.257 | 0.557 | 0.116 | 0.895 | 0.949 |
| Q17473 | 0.642 | 0.209 | 0.508 | 0.855 | 0.107 | 0.348 |
| Q17474 | 1.01 | 0.013 | 0.151 | 0.373 | 0.268 | 0.541 |
| Q17489 | -0.197 | 0.461 | 0.698 | 0.596 | 0.0482 | 0.26 |
| Q17490 | 0.335 | 0.51 | 0.734 | -0.272 | 0.591 | 0.794 |
| Q17571 | 0.947 | 0.0698 | 0.299 | 0.0974 | 0.834 | 0.92 |
| Q17686 | -1.71 | 0.126 | 0.391 | -1.17 | 0.274 | 0.548 |
| Q17698 | -1.18 | 0.0848 | 0.328 | -2.56 | 0.003 | 0.0849 |
| Q17763 | -0.251 | 0.219 | 0.516 | 0.0825 | 0.672 | 0.839 |
| Q17832 | -0.186 | 0.346 | 0.636 | -0.148 | 0.448 | 0.699 |
| Q17849 | -0.07 | 0.841 | 0.924 | -0.291 | 0.413 | 0.672 |
| Q17880 | 0.261 | 0.692 | 0.849 | -1.02 | 0.146 | 0.4 |
| Q17935 | 0.267 | 0.39 | 0.661 | 0.52 | 0.116 | 0.36 |
| Q17993 | 1.06 | 0.134 | 0.401 | 1.63 | 0.0342 | 0.232 |
| Q17994 | 0.684 | 0.0542 | 0.268 | 0.223 | 0.481 | 0.717 |
| Q18032 | 0.924 | 0.00198 | 0.0802 | 0.27 | 0.216 | 0.477 |
| Q18036 | 0.287 | 0.622 | 0.812 | 1.16 | 0.0741 | 0.311 |
| Q18074 | 1.5 | 0.00231 | 0.0825 | 1.59 | 0.00164 | 0.0731 |
| Q18100 | 1.31 | 0.0241 | 0.2 | 1.37 | 0.0199 | 0.191 |
| Q18124 | 0.836 | 0.131 | 0.396 | 0.281 | 0.586 | 0.791 |
| Q18231 | -0.364 | 0.137 | 0.406 | -0.313 | 0.193 | 0.451 |
| Q18265 | 1.11 | 0.355 | 0.638 | 2.86 | 0.0362 | 0.233 |
| Q18276 | 0.0823 | 0.85 | 0.931 | 0.25 | 0.571 | 0.783 |
| Q18577 | 0.214 | 0.458 | 0.697 | -0.129 | 0.65 | 0.831 |
| Q18599 | 1.02 | 0.00184 | 0.0798 | 0.707 | 0.0128 | 0.153 |
| Q18625 | 1.92 | 0.125 | 0.387 | -0.00317 | 0.998 | 0.998 |
| Q18853 | 1.05 | 0.132 | 0.397 | 0.86 | 0.208 | 0.47 |
| Q18886 | -1.05 | 0.397 | 0.661 | -0.821 | 0.504 | 0.737 |
| Q19007 | 0.498 | 0.155 | 0.434 | 1.01 | 0.0135 | 0.157 |
| Q19057 | 1.26 | 0.0328 | 0.214 | 1.17 | 0.0441 | 0.251 |
| Q19058 | 0.428 | 0.156 | 0.434 | -0.17 | 0.549 | 0.766 |
| Q19063 | 0.805 | 0.298 | 0.601 | 0.145 | 0.845 | 0.925 |
| Q19064 | -0.858 | 0.333 | 0.629 | -0.35 | 0.686 | 0.847 |
| Q19102 | -0.589 | 0.0198 | 0.186 | -0.357 | 0.114 | 0.36 |
| Q19240 | 1.43 | 0.228 | 0.527 | 0.819 | 0.474 | 0.712 |
| Q19246 | -3.33 | 0.03 | 0.212 | -3.21 | 0.0346 | 0.232 |
| Q19324 | -0.398 | 0.565 | 0.775 | 0.216 | 0.754 | 0.882 |
| Q19328 | 0.294 | 0.138 | 0.408 | -0.146 | 0.435 | 0.686 |
| Q19437 | 0.0343 | 0.889 | 0.941 | -0.197 | 0.431 | 0.686 |
| Q19478 | 2.3 | 0.0727 | 0.302 | -0.214 | 0.851 | 0.928 |
| Q19554 | -0.285 | 0.697 | 0.852 | -0.124 | 0.865 | 0.933 |
| Q19591 | -1.09 | 0.385 | 0.659 | -0.927 | 0.455 | 0.703 |
| Q19694 | 1.06 | 0.322 | 0.616 | 0.261 | 0.802 | 0.902 |
| Q19723 | 0.153 | 0.812 | 0.911 | 0.039 | 0.952 | 0.983 |
| Q19790 | 1.28 | 0.303 | 0.603 | 2.95 | 0.0353 | 0.232 |
| Q19973 | 2.19 | 0.0449 | 0.236 | 0.657 | 0.494 | 0.73 |
| Q1XFY9 | -0.24 | 0.321 | 0.616 | 0.245 | 0.311 | 0.59 |
| Q20011 | -1.09 | 0.0101 | 0.137 | -0.123 | 0.712 | 0.859 |
| Q20034 | -0.2 | 0.879 | 0.94 | 2.88 | 0.0542 | 0.277 |
| Q20049 | -0.904 | 0.429 | 0.681 | -0.613 | 0.588 | 0.791 |
| Q20107 | -1.09 | 0.192 | 0.481 | -0.91 | 0.269 | 0.541 |
| Q20122 | -0.507 | 0.705 | 0.855 | 1.63 | 0.243 | 0.504 |
| Q20173 | -0.678 | 0.635 | 0.819 | 0.272 | 0.848 | 0.926 |
| Q20203 | 0.62 | 0.0618 | 0.287 | 0.663 | 0.0489 | 0.263 |
| Q20206 | 0.14 | 0.411 | 0.67 | 0.3 | 0.102 | 0.339 |
| Q20239 | -0.474 | 0.168 | 0.448 | -0.473 | 0.169 | 0.421 |
| Q20264 | 1.35 | 0.217 | 0.514 | -0.666 | 0.527 | 0.746 |
| Q20276 | -0.548 | 0.347 | 0.637 | -1.51 | 0.0257 | 0.216 |
| Q20277 | -0.63 | 0.0309 | 0.212 | -0.182 | 0.469 | 0.711 |
| Q20306 | 3.03 | 0.0906 | 0.336 | 0.531 | 0.743 | 0.879 |
| Q20310 | -0.567 | 0.258 | 0.558 | -0.603 | 0.231 | 0.494 |
| Q20330 | 0.252 | 0.802 | 0.909 | 0.337 | 0.738 | 0.878 |
| Q20384 | -1.27 | 0.131 | 0.396 | -2.62 | 0.00876 | 0.134 |
| Q20546 | 0.813 | 0.395 | 0.661 | 0.246 | 0.792 | 0.9 |
| Q20603 | 1.36 | 0.0672 | 0.294 | 1.47 | 0.0516 | 0.269 |
| Q20634 | -0.0961 | 0.727 | 0.869 | 0.511 | 0.0925 | 0.33 |
| Q20637 | 0.385 | 0.0902 | 0.336 | 0.0363 | 0.86 | 0.931 |
| Q20660 | -2.29 | 0.147 | 0.422 | -0.521 | 0.723 | 0.868 |
| Q20676 | 1.05 | 0.0022 | 0.0825 | 0.794 | 0.00988 | 0.139 |
| Q20684 | 0.639 | 0.0193 | 0.186 | 0.405 | 0.0997 | 0.337 |
| Q20693 | 1.72 | 0.349 | 0.637 | 1.57 | 0.391 | 0.653 |
| Q20753 | -0.962 | 0.434 | 0.683 | -0.904 | 0.461 | 0.705 |
| Q20770 | -0.12 | 0.903 | 0.95 | 0.745 | 0.459 | 0.705 |
| Q20774 | 0.18 | 0.639 | 0.821 | 0.27 | 0.486 | 0.721 |
| Q20780 | 0.277 | 0.222 | 0.521 | 0.221 | 0.322 | 0.598 |
| Q20877 | -1.14 | 0.29 | 0.591 | 0.401 | 0.7 | 0.854 |
| Q20921 | -1.39 | 0.0246 | 0.2 | -0.72 | 0.188 | 0.448 |
| Q20950 | -0.0831 | 0.721 | 0.868 | -0.247 | 0.304 | 0.584 |
| Q21000 | 0.455 | 0.388 | 0.66 | -0.804 | 0.146 | 0.4 |
| Q21004 | 0.0994 | 0.862 | 0.936 | 1.32 | 0.0458 | 0.256 |
| Q21021 | -0.0295 | 0.986 | 0.994 | -0.206 | 0.9 | 0.95 |
| Q21233 | 0.792 | 0.175 | 0.458 | -0.291 | 0.598 | 0.798 |
| Q21274 | 0.166 | 0.84 | 0.924 | -1.49 | 0.0992 | 0.337 |
| Q21284 | -0.0364 | 0.969 | 0.984 | 0.121 | 0.898 | 0.95 |
| Q21342 | 0.961 | 0.215 | 0.514 | -0.0322 | 0.965 | 0.986 |
| Q21465 | 0.0493 | 0.802 | 0.909 | 0.172 | 0.394 | 0.654 |
| Q21481 | 0.964 | 0.255 | 0.554 | 1.33 | 0.13 | 0.379 |
| Q21525 | 0.754 | 0.107 | 0.361 | 0.859 | 0.073 | 0.309 |
| Q21544 | 0.785 | 0.446 | 0.692 | -0.499 | 0.624 | 0.813 |
| Q21559 | 0.145 | 0.497 | 0.724 | -0.372 | 0.108 | 0.35 |
| Q21732 | 0.132 | 0.719 | 0.867 | -0.335 | 0.37 | 0.641 |
| Q21742 | 0.28 | 0.815 | 0.912 | -0.998 | 0.414 | 0.672 |
| Q21746 | -0.511 | 0.0831 | 0.324 | -0.102 | 0.703 | 0.854 |
| Q21763 | 0.149 | 0.677 | 0.843 | 0.292 | 0.422 | 0.68 |
| Q21888 | -0.11 | 0.929 | 0.965 | 2.69 | 0.0548 | 0.277 |
| Q21938 | -3.35 | 0.00524 | 0.107 | -2.96 | 0.00979 | 0.139 |
| Q22101 | -0.192 | 0.425 | 0.679 | -0.151 | 0.526 | 0.746 |
| Q22135 | 1.19 | 0.193 | 0.481 | -0.655 | 0.457 | 0.703 |
| Q22291 | -0.801 | 0.549 | 0.765 | -0.916 | 0.495 | 0.73 |
| Q22352 | -0.658 | 0.372 | 0.654 | -1.78 | 0.0348 | 0.232 |
| Q22370 | 0.324 | 0.179 | 0.463 | 1.01 | 0.00193 | 0.0743 |
| Q22442 | -0.38 | 0.217 | 0.514 | -0.293 | 0.33 | 0.608 |
| Q22483 | 0.0939 | 0.886 | 0.941 | 0.163 | 0.803 | 0.902 |
| Q22508 | 0.727 | 0.0362 | 0.221 | 0.0113 | 0.97 | 0.986 |
| Q22515 | -0.567 | 0.568 | 0.775 | 2.64 | 0.0251 | 0.215 |
| Q22562 | -0.201 | 0.482 | 0.713 | -0.59 | 0.0633 | 0.294 |
| Q22615 | -0.454 | 0.639 | 0.821 | -0.842 | 0.393 | 0.653 |
| Q22620 | 1.26 | 0.354 | 0.638 | -0.0325 | 0.98 | 0.989 |
| Q22666 | -1.67 | 0.0275 | 0.209 | -0.0201 | 0.975 | 0.986 |
| Q22714 | 2.33 | 0.0166 | 0.171 | 2.84 | 0.00634 | 0.117 |
| Q22716 | -0.304 | 0.114 | 0.371 | -0.161 | 0.373 | 0.642 |
| Q22719 | 0.204 | 0.433 | 0.683 | 0.632 | 0.0349 | 0.232 |
| Q22768 | 0.121 | 0.872 | 0.939 | 0.536 | 0.483 | 0.718 |
| Q22781 | -0.445 | 0.492 | 0.721 | 0.28 | 0.662 | 0.837 |
| Q22850 | 1.45 | 0.0126 | 0.151 | 1.36 | 0.0174 | 0.186 |
| Q22968 | 0.547 | 0.146 | 0.422 | 0.244 | 0.492 | 0.728 |
| Q23050 | -0.843 | 0.0228 | 0.196 | -0.802 | 0.028 | 0.217 |
| Q23098 | 2.49 | 0.141 | 0.413 | 2.31 | 0.168 | 0.42 |
| Q23186 | -0.268 | 0.726 | 0.869 | -0.115 | 0.879 | 0.942 |
| Q23258 | 2.05 | 0.0789 | 0.314 | 1.74 | 0.124 | 0.372 |
| Q23315 | 0.487 | 0.467 | 0.699 | -0.579 | 0.391 | 0.653 |
| Q23330 | -0.198 | 0.571 | 0.778 | 0.882 | 0.0312 | 0.226 |
| Q23451 | -0.581 | 0.161 | 0.437 | 0.0531 | 0.891 | 0.948 |
| Q23487 | -0.164 | 0.82 | 0.916 | -0.901 | 0.234 | 0.497 |
| Q23543 | 0.0558 | 0.923 | 0.963 | -0.468 | 0.429 | 0.685 |
| Q23597 | 1.01 | 0.00455 | 0.107 | 0.535 | 0.0698 | 0.302 |
| Q23621 | 0.103 | 0.575 | 0.78 | -0.00602 | 0.973 | 0.986 |
| Q23624 | 1.65 | 0.0105 | 0.138 | 0.461 | 0.373 | 0.642 |
| Q23642 | 1.13 | 0.241 | 0.54 | -0.467 | 0.615 | 0.807 |
| Q2EEM8 | 1.42 | 0.486 | 0.717 | -0.584 | 0.771 | 0.89 |
| Q2XN18 | -0.351 | 0.284 | 0.586 | 0.488 | 0.15 | 0.402 |
| Q3LFN1 | 0.434 | 0.453 | 0.696 | 0.0372 | 0.948 | 0.983 |
| Q45EK1 | -0.558 | 0.513 | 0.736 | -0.0249 | 0.976 | 0.986 |
| Q5FC71 | -0.249 | 0.426 | 0.68 | -0.956 | 0.0129 | 0.153 |
| Q5FC82 | -0.0777 | 0.904 | 0.95 | 1.72 | 0.0261 | 0.216 |
| Q65XX1 | 0.262 | 0.593 | 0.793 | 0.171 | 0.726 | 0.871 |
| Q65XX4 | -1.68 | 0.111 | 0.367 | -1.6 | 0.125 | 0.372 |
| Q69Z13 | 2.33 | 0.0812 | 0.32 | 1.6 | 0.206 | 0.47 |
| Q6A575 | 0.806 | 0.454 | 0.696 | 0.39 | 0.714 | 0.859 |
| Q75MI6 | -1.32 | 0.396 | 0.661 | -2.36 | 0.148 | 0.4 |
| Q7JPE4 | 0.369 | 0.453 | 0.696 | -0.141 | 0.77 | 0.89 |
| Q7Z072 | -0.0414 | 0.861 | 0.936 | 0.158 | 0.51 | 0.741 |
| Q7Z1Q3 | -0.0179 | 0.972 | 0.985 | 0.106 | 0.837 | 0.921 |
| Q7Z2A9 | -2.33 | 0.0152 | 0.163 | -1.69 | 0.055 | 0.277 |
| Q86B36 | -1.24 | 0.166 | 0.446 | -0.863 | 0.32 | 0.598 |
| Q86FL8 | -0.77 | 0.185 | 0.47 | -1.4 | 0.0306 | 0.224 |
| Q86LS4 | -1.2 | 0.00754 | 0.121 | -0.639 | 0.0926 | 0.33 |
| Q86NC1 | 0.236 | 0.279 | 0.583 | 0.154 | 0.469 | 0.711 |
| Q86NC2 | -0.00182 | 0.997 | 0.998 | 0.352 | 0.526 | 0.746 |
| Q86NE0 | 0.96 | 0.271 | 0.573 | 0.481 | 0.57 | 0.782 |
| Q86NH9 | 1.05 | 0.099 | 0.349 | 1.13 | 0.0805 | 0.312 |
| Q8MPX7 | 1.02 | 0.0839 | 0.325 | 0.719 | 0.201 | 0.462 |
| Q8MXD9 | -0.00154 | 0.997 | 0.998 | -1.06 | 0.0348 | 0.232 |
| Q8MXH3 | -1.17 | 0.149 | 0.424 | -1.01 | 0.207 | 0.47 |
| Q8MXH7 | -0.887 | 0.204 | 0.499 | -1.53 | 0.0456 | 0.256 |
| Q8MXI1 | -0.628 | 0.456 | 0.697 | -0.778 | 0.361 | 0.632 |
| Q8MXR2 | 0.662 | 0.543 | 0.762 | 0.976 | 0.376 | 0.645 |
| Q8MXR6 | -0.161 | 0.653 | 0.828 | 0.602 | 0.12 | 0.367 |
| Q8T7Z3 | -1.15 | 0.00585 | 0.109 | -0.0562 | 0.858 | 0.931 |
| Q8WQA8 | 0.104 | 0.57 | 0.778 | -0.0339 | 0.852 | 0.928 |
| Q93168 | 0.244 | 0.485 | 0.716 | 0.103 | 0.765 | 0.887 |
| Q93204 | 0.857 | 0.213 | 0.514 | 2.5 | 0.00463 | 0.0997 |
| Q93315 | 1.46 | 0.0325 | 0.214 | 0.236 | 0.685 | 0.847 |
| Q93576 | -0.83 | 0.00805 | 0.121 | -0.668 | 0.0222 | 0.202 |
| Q93637 | -0.661 | 0.0247 | 0.2 | -0.343 | 0.188 | 0.448 |
| Q93791 | 1.3 | 0.00169 | 0.0762 | 0.813 | 0.0194 | 0.189 |
| Q93805 | 1.99 | 0.253 | 0.552 | 1.6 | 0.348 | 0.622 |
| Q93831 | 0.253 | 0.37 | 0.652 | 0.533 | 0.0815 | 0.312 |
| Q93838 | 0.681 | 0.545 | 0.762 | -2.01 | 0.1 | 0.338 |
| Q93871 | 0.264 | 0.769 | 0.893 | 1.34 | 0.163 | 0.416 |
| Q93896 | -1.88 | 0.079 | 0.314 | -2.05 | 0.0599 | 0.291 |
| Q93934 | -0.43 | 0.11 | 0.364 | -0.515 | 0.0633 | 0.294 |
| Q94055 | -0.387 | 0.693 | 0.849 | 0.338 | 0.73 | 0.873 |
| Q94162 | -0.0642 | 0.956 | 0.977 | 0.315 | 0.785 | 0.896 |
| Q94246 | 0.195 | 0.338 | 0.632 | 0.124 | 0.534 | 0.751 |
| Q94269 | 0.0553 | 0.886 | 0.941 | 0.0289 | 0.94 | 0.977 |
| Q94271 | -2.32 | 0.00514 | 0.107 | -1.61 | 0.029 | 0.22 |
| Q95PW6 | 0.18 | 0.699 | 0.853 | 0.0186 | 0.968 | 0.986 |
| Q95PW9 | 1.21 | 0.194 | 0.482 | 1.42 | 0.134 | 0.384 |
| Q95PX7 | -0.077 | 0.887 | 0.941 | -0.806 | 0.166 | 0.418 |
| Q95PZ1 | 0.362 | 0.615 | 0.809 | -1.85 | 0.0293 | 0.221 |
| Q95QC2 | 2.11 | 0.078 | 0.314 | 1.64 | 0.154 | 0.407 |
| Q95QS3 | -0.375 | 0.606 | 0.804 | -0.479 | 0.512 | 0.742 |
| Q95XD4 | 0.0103 | 0.993 | 0.998 | -0.359 | 0.773 | 0.89 |
| Q95XG6 | -2.15 | 0.0585 | 0.278 | -1.3 | 0.219 | 0.48 |
| Q95XJ0 | 0.507 | 0.0232 | 0.199 | 0.0172 | 0.926 | 0.965 |
| Q95XN6 | 0.648 | 0.44 | 0.685 | 0.177 | 0.829 | 0.917 |
| Q95XR0 | 0.812 | 0.0201 | 0.186 | 0.747 | 0.0285 | 0.218 |
| Q95XR1 | 0.529 | 0.157 | 0.434 | 1.32 | 0.00493 | 0.1 |
| Q95XS1 | -2.06 | 0.178 | 0.461 | -1.14 | 0.434 | 0.686 |
| Q95XS2 | 0.152 | 0.586 | 0.789 | -0.396 | 0.177 | 0.434 |
| Q95XT5 | -0.457 | 0.205 | 0.501 | -0.546 | 0.139 | 0.387 |
| Q95Y29 | 2.35 | 0.0534 | 0.266 | 4.14 | 0.00427 | 0.0992 |
| Q95Y87 | -0.234 | 0.787 | 0.905 | 1.58 | 0.0971 | 0.336 |
| Q95Y93 | -0.265 | 0.691 | 0.849 | 2.2 | 0.0098 | 0.139 |
| Q95YC6 | 0.193 | 0.493 | 0.721 | 0.579 | 0.0648 | 0.297 |
| Q95YC7 | 1.15 | 0.0501 | 0.253 | -0.217 | 0.673 | 0.839 |
| Q95YD5 | 0.683 | 0.619 | 0.811 | 0.419 | 0.759 | 0.886 |
| Q95YE7 | 0.273 | 0.752 | 0.88 | 1.43 | 0.127 | 0.372 |
| Q95ZL1 | 0.114 | 0.66 | 0.831 | -0.276 | 0.305 | 0.584 |
| Q95ZS5 | -1.57 | 0.0244 | 0.2 | -1 | 0.113 | 0.358 |
| Q95ZY7 | -0.457 | 0.465 | 0.699 | 0.213 | 0.73 | 0.873 |
| Q965Q1 | -0.894 | 0.0202 | 0.186 | -0.857 | 0.0243 | 0.211 |
| Q965V4 | 3.26 | 0.161 | 0.438 | 3.5 | 0.136 | 0.385 |
| Q966C7 | -0.343 | 0.522 | 0.743 | -0.147 | 0.781 | 0.896 |
| Q966I7 | -0.358 | 0.265 | 0.566 | -0.0307 | 0.92 | 0.964 |
| Q966I8 | 0.495 | 0.248 | 0.545 | 0.46 | 0.279 | 0.553 |
| Q967F1 | 0.922 | 0.411 | 0.67 | 1.83 | 0.125 | 0.372 |
| Q9BI71 | 0.323 | 0.67 | 0.839 | 2.42 | 0.0114 | 0.146 |
| Q9BIB7 | 0.48 | 0.739 | 0.875 | 1.51 | 0.31 | 0.588 |
| Q9BIC3 | 0.612 | 0.0387 | 0.226 | -0.245 | 0.349 | 0.622 |
| Q9BKP8 | -0.383 | 0.28 | 0.583 | -0.799 | 0.043 | 0.249 |
| Q9BKQ9 | -0.555 | 0.266 | 0.568 | -0.746 | 0.147 | 0.4 |
| Q9BKU5 | 0.345 | 0.109 | 0.363 | 0.222 | 0.278 | 0.551 |
| Q9BL03 | -0.39 | 0.222 | 0.521 | -0.293 | 0.349 | 0.622 |
| Q9BL27 | 0.72 | 0.204 | 0.499 | 1.03 | 0.083 | 0.316 |
| Q9BL33 | 1.7 | 0.0337 | 0.217 | 1.32 | 0.0812 | 0.312 |
| Q9BL34 | 0.311 | 0.365 | 0.648 | 0.519 | 0.148 | 0.4 |
| Q9BL39 | -0.202 | 0.863 | 0.936 | -0.878 | 0.46 | 0.705 |
| Q9BL61 | -0.649 | 0.466 | 0.699 | -0.757 | 0.399 | 0.659 |
| Q9GPA1 | 0.151 | 0.877 | 0.94 | -0.638 | 0.518 | 0.744 |
| Q9GRY9 | 0.523 | 0.0877 | 0.332 | 0.614 | 0.0522 | 0.27 |
| Q9GRZ0 | -1.38 | 0.184 | 0.469 | -1.96 | 0.073 | 0.309 |
| Q9GRZ9 | 0.443 | 0.237 | 0.538 | 0.382 | 0.302 | 0.584 |
| Q9GYI1 | 0.455 | 0.475 | 0.708 | 0.205 | 0.744 | 0.879 |
| Q9GYQ6 | -2.09 | 0.044 | 0.232 | 0.229 | 0.798 | 0.902 |
| Q9GZH5 | -0.823 | 0.329 | 0.623 | -1.52 | 0.0924 | 0.33 |
| Q9N362 | -2.95 | 0.000929 | 0.0538 | -2.23 | 0.00467 | 0.0997 |
| Q9N384 | -0.242 | 0.571 | 0.778 | -0.906 | 0.0592 | 0.289 |
| Q9N3D9 | -1.63 | 0.00315 | 0.0912 | -1.37 | 0.00801 | 0.132 |
| Q9N3F7 | 0.487 | 0.296 | 0.6 | 0.191 | 0.672 | 0.839 |
| Q9N3G0 | 0.0617 | 0.858 | 0.936 | 0.267 | 0.446 | 0.696 |
| Q9N3H3 | 0.26 | 0.503 | 0.729 | -0.0486 | 0.899 | 0.95 |
| Q9N3T3 | 1.19 | 0.0264 | 0.203 | 0.817 | 0.0968 | 0.336 |
| Q9N456 | -0.466 | 0.253 | 0.552 | 0.766 | 0.0787 | 0.312 |
| Q9N492 | 0.0261 | 0.977 | 0.989 | 1.66 | 0.0908 | 0.329 |
| Q9N4C6 | 1.12 | 0.107 | 0.361 | 1.26 | 0.0754 | 0.311 |
| Q9N4F3 | 2.43 | 0.00498 | 0.107 | 0.277 | 0.668 | 0.839 |
| Q9N4G8 | 0.404 | 0.625 | 0.813 | -0.187 | 0.82 | 0.914 |
| Q9N4H7 | 1.86 | 0.0285 | 0.212 | -1.18 | 0.126 | 0.372 |
| Q9N4I3 | -0.275 | 0.318 | 0.613 | -0.272 | 0.324 | 0.6 |
| Q9N4L8 | -1.24 | 0.0406 | 0.229 | -0.337 | 0.524 | 0.746 |
| Q9N4Y8 | 0.401 | 0.159 | 0.434 | 0.275 | 0.316 | 0.594 |
| Q9N5B3 | 0.119 | 0.659 | 0.831 | 0.348 | 0.218 | 0.48 |
| Q9N5E4 | 0.0392 | 0.861 | 0.936 | -0.0403 | 0.857 | 0.931 |
| Q9N5S7 | -0.00013 | 1 | 1 | 0.451 | 0.281 | 0.557 |
| Q9N5V3 | -1.27 | 0.066 | 0.293 | -0.169 | 0.783 | 0.896 |
| Q9NA32 | 1.87 | 0.0292 | 0.212 | 1.8 | 0.0339 | 0.232 |
| Q9NA39 | 0.521 | 0.0405 | 0.229 | 0.426 | 0.0809 | 0.312 |
| Q9NA98 | 0.00849 | 0.994 | 0.998 | 0.0324 | 0.976 | 0.986 |
| Q9NAB2 | 0.923 | 0.157 | 0.434 | 1.76 | 0.0186 | 0.188 |
| Q9NAI5 | -0.408 | 0.351 | 0.637 | 0.205 | 0.631 | 0.819 |
| Q9NES7 | 1.55 | 0.00135 | 0.0656 | 0.609 | 0.0908 | 0.329 |
| Q9NEY7 | 0.4 | 0.548 | 0.765 | 0.383 | 0.565 | 0.78 |
| Q9NLD1 | 0.674 | 0.0248 | 0.2 | 0.571 | 0.0474 | 0.259 |
| Q9TXI4 | -0.134 | 0.886 | 0.941 | -0.744 | 0.434 | 0.686 |
| Q9TXU7 | -0.765 | 0.0956 | 0.345 | -0.38 | 0.374 | 0.643 |
| Q9TXY8 | -0.955 | 0.445 | 0.691 | 0.206 | 0.866 | 0.933 |
| Q9TYS3 | -0.674 | 0.0314 | 0.212 | -0.473 | 0.104 | 0.343 |
| Q9TYX1 | 0.361 | 0.782 | 0.903 | -0.658 | 0.615 | 0.807 |
| Q9TZ68 | 0.709 | 0.0987 | 0.349 | 0.588 | 0.16 | 0.41 |
| Q9TZS5 | -0.0379 | 0.918 | 0.958 | 0.883 | 0.0398 | 0.237 |
| Q9U1Q1 | -0.804 | 0.4 | 0.662 | 1.66 | 0.104 | 0.344 |
| Q9U1X9 | -2.37 | 0.00651 | 0.116 | -0.962 | 0.173 | 0.427 |
| Q9U228 | 0.128 | 0.93 | 0.966 | 4.4 | 0.0154 | 0.17 |
| Q9U229 | -0.244 | 0.565 | 0.775 | -0.424 | 0.328 | 0.606 |
| Q9U238 | -0.173 | 0.82 | 0.916 | 0.63 | 0.419 | 0.679 |
| Q9U296 | 1.47 | 0.0777 | 0.314 | 0.00612 | 0.993 | 0.997 |
| Q9U2Q8 | -0.642 | 0.00849 | 0.125 | -0.815 | 0.00237 | 0.0758 |
| Q9U2U0 | 0.324 | 0.778 | 0.9 | 0.678 | 0.559 | 0.777 |
| Q9U302 | -0.127 | 0.458 | 0.697 | -0.337 | 0.0728 | 0.309 |
| Q9U307 | 0.603 | 0.489 | 0.719 | 0.736 | 0.403 | 0.661 |
| Q9U315 | -1.28 | 0.164 | 0.443 | -0.752 | 0.393 | 0.653 |
| Q9U329 | 0.287 | 0.341 | 0.634 | 0.312 | 0.304 | 0.584 |
| Q9U382 | 2.42 | 0.00478 | 0.107 | 0.682 | 0.302 | 0.584 |
| Q9U3H4 | 0.708 | 0.0294 | 0.212 | 0.117 | 0.669 | 0.839 |
| Q9U3P5 | 1.02 | 0.337 | 0.632 | 0.127 | 0.901 | 0.95 |
| Q9U9J8 | 0.956 | 0.00227 | 0.0825 | 0.954 | 0.0023 | 0.0758 |
| Q9UAQ6 | 0.405 | 0.0717 | 0.3 | -0.0726 | 0.718 | 0.863 |
| Q9UAY9 | 1.37 | 0.308 | 0.608 | 2.38 | 0.0967 | 0.336 |
| Q9UAZ2 | -2.23 | 0.02 | 0.186 | -2.28 | 0.0179 | 0.186 |
| Q9UB28 | 0.28 | 0.836 | 0.923 | -1.42 | 0.31 | 0.588 |
| Q9XTV4 | -0.318 | 0.275 | 0.577 | -0.00982 | 0.972 | 0.986 |
| Q9XTY9 | 0.701 | 0.584 | 0.787 | -0.945 | 0.464 | 0.709 |
| Q9XTZ2 | -1.04 | 0.346 | 0.636 | -1.48 | 0.19 | 0.448 |
| Q9XU56 | -2.44 | 0.0348 | 0.218 | -1.46 | 0.164 | 0.416 |
| Q9XU97 | 0.612 | 0.0306 | 0.212 | 0.535 | 0.0508 | 0.268 |
| Q9XUE5 | 1.24 | 0.214 | 0.514 | 1.53 | 0.133 | 0.384 |
| Q9XUL7 | 1.47 | 0.421 | 0.676 | 2.24 | 0.235 | 0.497 |
| Q9XUT0 | 1.83 | 0.0788 | 0.314 | -0.284 | 0.761 | 0.886 |
| Q9XUW5 | 0.0395 | 0.969 | 0.984 | -0.456 | 0.658 | 0.836 |
| Q9XVE9 | -0.707 | 0.0465 | 0.24 | -0.399 | 0.218 | 0.48 |
| Q9XVJ3 | -1.2 | 0.00276 | 0.0841 | -0.483 | 0.119 | 0.366 |
| Q9XVQ2 | 0.483 | 0.397 | 0.661 | -1.11 | 0.0758 | 0.311 |
| Q9XW20 | 0.899 | 0.614 | 0.809 | -0.23 | 0.896 | 0.949 |
| Q9XW37 | 0.11 | 0.768 | 0.893 | 0.459 | 0.239 | 0.503 |
| Q9XW41 | 0.498 | 0.172 | 0.455 | 0.866 | 0.0321 | 0.228 |
| Q9XWG2 | 0.581 | 0.101 | 0.349 | 1.03 | 0.0119 | 0.151 |
| Q9XWJ5 | 0.817 | 0.202 | 0.498 | 0.179 | 0.767 | 0.889 |
| Q9XWL1 | -0.721 | 0.296 | 0.599 | -1.71 | 0.0297 | 0.223 |
| Q9XWN4 | -1.83 | 0.128 | 0.395 | -2.66 | 0.0396 | 0.237 |
| Q9XWP7 | -0.403 | 0.0902 | 0.336 | -0.343 | 0.138 | 0.387 |
| Q9XWS4 | -0.394 | 0.153 | 0.431 | -0.238 | 0.367 | 0.637 |
| Q9XWS6 | 3.58 | 0.00376 | 0.101 | 2.07 | 0.046 | 0.256 |
| Q9XWT3 | -0.443 | 0.69 | 0.849 | 0.104 | 0.925 | 0.965 |
| Q9XWU9 | 0.333 | 0.362 | 0.645 | 0.255 | 0.48 | 0.716 |
| Q9XX57 | -1.01 | 0.0145 | 0.16 | -1.04 | 0.0123 | 0.153 |
| Q9XXE2 | -1.98 | 0.0568 | 0.273 | -2.4 | 0.0276 | 0.217 |
| Q9XXR4 | -3.47 | 0.0549 | 0.27 | -3.02 | 0.0855 | 0.32 |
| Q9XXR5 | -1.01 | 0.0944 | 0.343 | -1.6 | 0.0176 | 0.186 |
| V6CKH6 | 0.454 | 0.276 | 0.579 | 0.694 | 0.112 | 0.358 |
